# Speed modulations in grid cell information geometry

**DOI:** 10.1101/2024.09.18.613797

**Authors:** Zeyuan Ye, Ralf Wessel

## Abstract

Grid cells, known for their hexagonal spatial firing patterns, are widely regarded as essential to the brain’s internal representation of the external space. Maintaining an accurate internal spatial representation is challenging when an animal is running at high speeds, as its self-location constantly changes. Previous studies of speed modulation of grid cells focused on individual or pairs of grid cells, yet neurons represent information via collective population activity. Population noise covariance can have significant impact on information coding that is impossible to infer from individual neuron analysis. To address this issue, we developed a novel Gaussian Process with Kernel Regression (GKR) method that allows study the simultaneously recorded neural population representation from an information geometry framework. We applied GKR to grid cell population activity, and found that running speed increases both grid cell activity toroidal-like manifold size and noise strength. Importantly, the effect of manifold dilation outpaces the effect of noise increasement, as indicated by the overall higher Fisher information at increasing speeds. This result is further supported by improved spatial information decoding accuracy at high speeds. Finally, we showed that the existence of noise covariance is information detrimental because it causes more noise projected onto the manifold surface. In total, our results indicate that grid cell spatial coding improves with increasing running speed. GKR provides a useful tool to understand neural population coding from an intuitive information geometric perspective.

## Introduction

In navigation, it is crucial that the brain forms certain internal representation of the external space^1^. Grid cells are widely regarded as an essential component of internal spatial representation^2,3^. Their hexagonal spatial firing patterns are thought to form a coordinate system of external space^4^ and support the downstream hippocampal spatial representations (e.g., place cells)^5–9^. However, establishing an accurate internal spatial coding is challenging, especially when the subject is running at high speed, where self-location constantly changes^10^. The effect of running speed modulation on grid cell population coding remains unclear.

Previous literature offers dual possible predictions about running speed modulation on grid cell codes. On the one hand, speed may support grid cell spatial coding. Running speed is known to mostly increase grid cell firing rates^11–13^. Rats running at a high speeds (10 cm/s to 50 cm/s) are also known to have more medial entorhinal cortex (MEC) cells coding spatial information than when running at a low speeds (2 cm/s to 10 cm/s)^10^. On the other hand, speed signals disrupt the phase differences between pairs of grid cells^14^. Increasing speed may also lead to larger input noise (possibly from medial septum^11,15^ or speed cells^16,17^), causing larger noise error to accumulate over time, thus degrading spatial coding fidelity^18–21^.

While these previous studies provide insights into speed modulations of grid cells, their analyses were limited to individual or pairs of grid cells^11–14^ (although decoding analysis has been performed on heterogeneous MEC cell population^10^). Neurons in the brain represent information through their collective population activity. Population noise covariance can significantly impact information coding, depending on its fine structure^22–28^. Grid cells’ activities are especially known to be tightly coupled and change coherently^14,29^. To study the speed modulation of grid cell code, it is important to analysis simultaneously recorded grid cell population activities, including the effect of noise covariance. Yet such a study is still lacking.

Inferring noise covariance is challenging due to the high-dimensional nature of neural data. When the information value is discretized, sample covariance is a common approach to infer noise covariance^23^. When the information is continuous, a practical approach is to first discretize the information values (e.g., orientations of static grating stimuli). Experimentalists then perform trial-based experiments on these discretized values^30–33^. Trial-based data allows for inferring noise covariance using sample covariance or, more recently, Wishart processes as implemented by Nejatbakhsh et al^34^.

However, in many cases, discretizing continuous information values and obtaining trial-based data is impractical: (1) the dimensionality of the information itself can be high, causing an exponential number of required discretized values (e.g., when the input is natural images^27^), and (2) some experiments, particularly naturalistic experiments, do not have repeated trials^31,33^ (e.g. navigation tasks^35^). A study on retinal representation of natural images circumvented these challenges by replacing retinal data with a convolutional neural network (CNN) unit, thereby explicitly formulating the noise covariance^27^. Yet, this approach is mainly based on the discovered remarkable similarities between retinal neurons and CNN units^36^. There’s a trending in neuroscience to move beyond trial-based experiments, towards trial-free naturalistic experiments^31,33^. However, to our knowledge in the broader field of neuroscience, there is no reliable method for inferring noise covariance from high-dimensional neural data in naturalistic tasks without repeated trials.

In this paper, we propose a novel method called Gaussian Process with Kernel Regression (GKR), which enables the inference of both the smooth mean (manifold) and noise covariance from high-dimensional neural data, including data from naturalistic tasks. The study of manifolds and noise covariance formally fall within the framework of information geometry^27^. We applied GKR to simultaneously recorded grid cell activities^35^. We found that: (1) Running speed both dilates the grid cells’ toroidal-like manifold and increases noise; (2) Nevertheless, the effect of manifold dilation outpaces the effect of noise increase, as indicated by the overall higher Fisher information at increasing speeds, and further supported by improved spatial coding accuracy at higher speeds; (3) Furthermore, compared to hypothetical independently firing grid cells, we found that noise correlations in real grid cells “reshape/orient” the noise structure such that more noise is projected onto the manifold surface, indicating that noise correlation in grid cells is information-detrimental. Overall, our results indicate that running speed enhances grid cell spatial coding through geometric modulations. GKR provides a powerful tool to interpret noisy neural data from an intuitive information geometry perspective.

## Results

### Grid cell population spatial coding accuracy improves with increasing speed

Grid cell population activities were recorded as rats performed open-field foraging (OF) tasks^35^. This yielded about 100–180 simultaneously recorded grid cells, varying by experimental configurations— specifically, the rat, recording day, and grid cell module (see Methods). The experiment resulted in a total of nine distinct experimental configurations^35^. For clarity in referencing, we employed a notation system; for instance, “R1M2” denotes data collected from rat “R” on day 1, specifically from grid cell module 2. Grid cells within the same module have similar spatial periods but different phases. We converted the raw spiking data into firing rates. Example grid cells’ rate maps are shown in Figure 1A and SI Figure 1.

**Figure 1.**
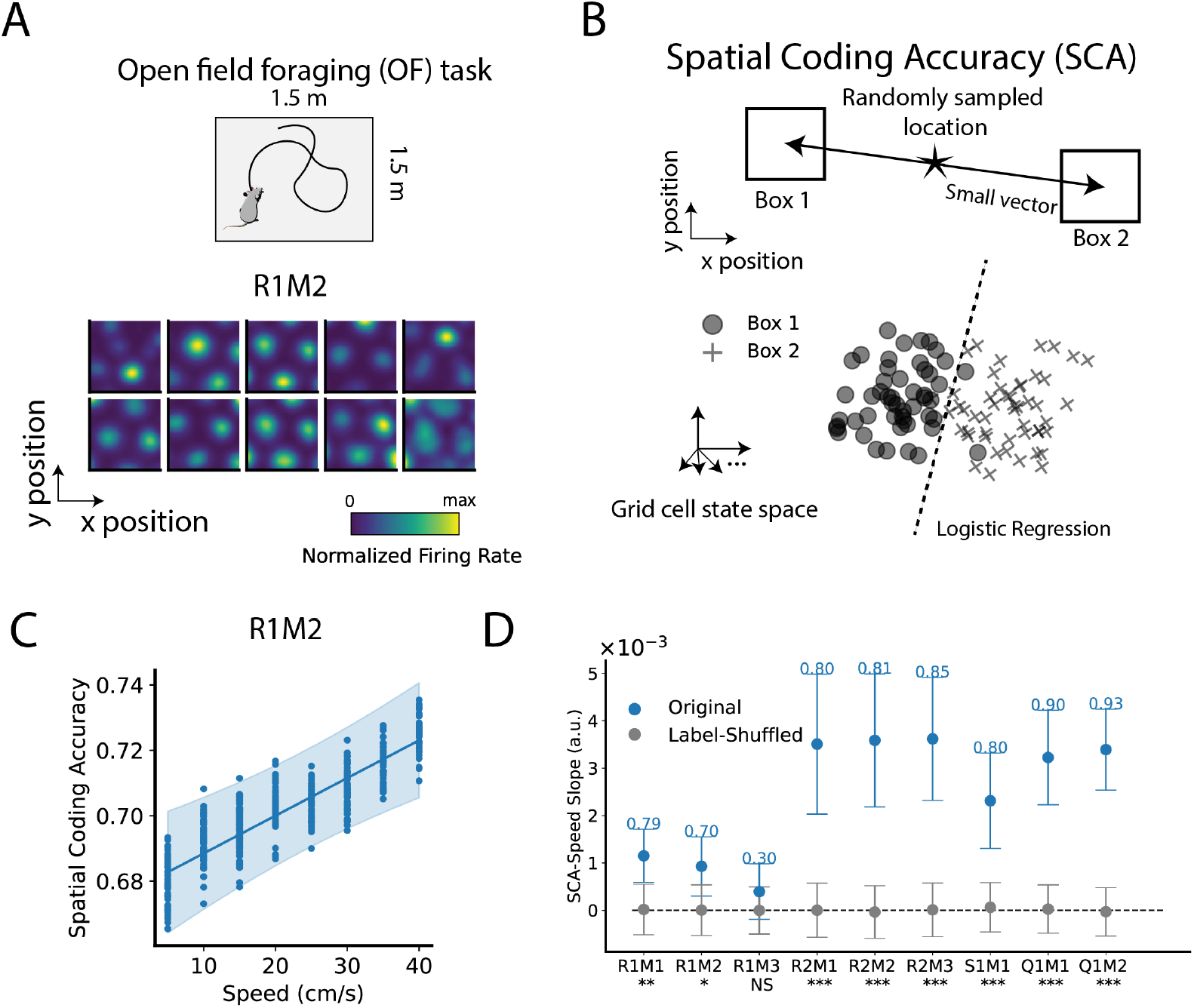
Grid cell population spatial coding accuracy (SCA) improves with increasing speed. (**A**) Top: Rat in an open-field foraging (OF) task with grid cell population activities (neural states for short) recorded. Nine experimental configurations (different rats, days and grid cell modules) were inspected. For example, experimental condition R1M2 represents rat ‘R’, day 1, grid cell module 2. Data in each experimental configuration were subsampled, generating 50 sampled datasets 𝒟_*s*_. 𝒟_*s*_ has similar number of data points at different speed value (see texts and Methods). Bottom: Example grid cells’ rate map in R1M2. The other examples can be found in SI Figure 1. (**B**) SCA measures the quality of grid cell spatial coding. Random locations were sampled in the OF. For each location, we drew two opposite but close boxes (separated by small vectors). Neural states in the two boxes were collected and classified by a logistic regression. The average classification accuracy across random sampled locations is the SCA (see Methods). (**C**) SCA as a function of rat’s speed. Each dot represents the SCA computed from a sampled dataset 𝒟_*s*_ at one speed bin (see Methods). Solid line and error band show the best-fitting line and 95% confidence interval (CI) using Bayesian linear ensemble averaging (BLEA, see texts and Methods). (**D**) SCA-speed slope of different experimental configurations. Dots and error bars represent mean and 95% CI of the slope. Numbers above error bars are linear models’ r-squared values (see Methods). Stars below x-axis labels indicate the significance level of whether the slope from original datasets 𝒟_*s*_ differs statistically from label-shuffled data (Bayesian method, see Methods). ***: p < 0.001; **: p < 0.01; *: p < 0.05; NS: no significance.

We analyzed the rats’ behavior data and found that rats tend to stay at low speeds, meaning there are more data points in the low-speed region than in the high-speed region (SI Figure 2). This can lead to future possible biased analysis. Therefore, we randomly sampled the data at different running speed bins, such that the sampled dataset 𝒟_*s*_ has a same number of data points in each speed bin (bin width = 5 cm/s, see Methods). Fifty 𝒟_*s*_ were sampled.

**Figure 2.**
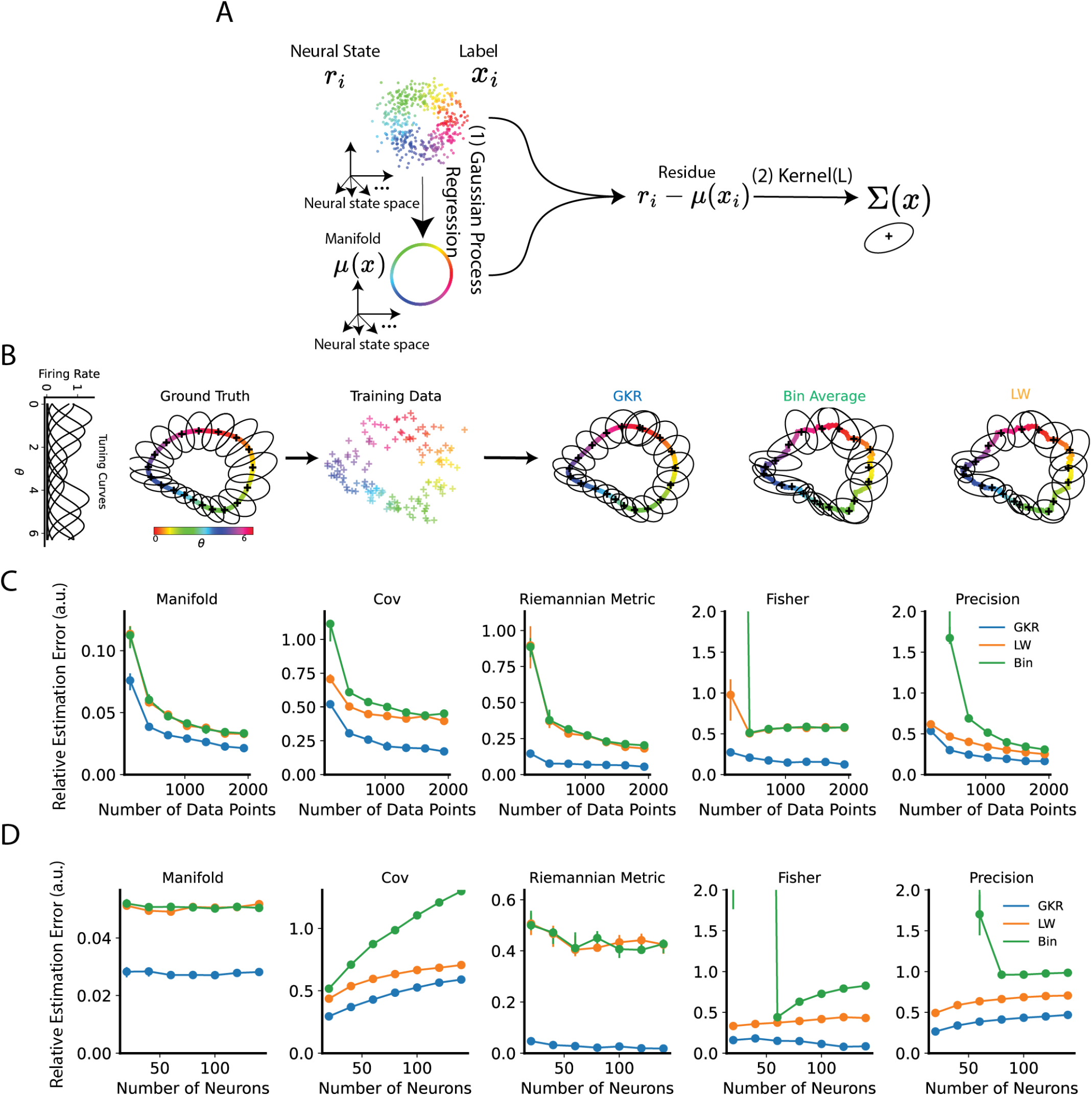
Gaussian Process with Kernel Regression (GKR) can infer the smooth manifold and covariance from noisy data. (**A**) Given neural states ***r*** and labels ***x*** (label can be stimulus parameters, animal behavior, animal position etc.), the goal of GKR is to infer the conditional distribution *p*(***r***|***x***) as a smooth function of ***x***. GKR has two major steps. In step one, Gaussian process regression is used to fit a mean function ***μ***(***x***) (i.e. manifold). In step two, using the residual at data points, i.e. ***r***_*&*_− ***μ***(***x***_*&*_), GKR uses a kernel method to estimate a smoothly varying noise covariance Σ(***x***). The resulting *p*(***r***|***x***) is approximated as a Gaussian distribution. (**B**) Applying GKR to a synthetic dataset. The synthetic dataset has *N* synthetic neurons with heterogeneous tuning curves to a circular label *θ* ranging from 0 to 2π. Ground truth ***μ***(*θ*) and Σ(*θ*) are visualized on the first two principal components plane (via PCA), shown on the left. Ellipses major axes represent the direction of eigenvectors with lengths proportional to eigenvalues. In this example, the synthetic data set has 10 neurons and generated 100 data points. These data points were used for fitting GKR, bin average, and Ledoit-Wolf methods (see methods), with fitting results shown on the right. (**C, D**) Quantifications of the estimation performances. The default number of data points is 300, and the default number of neurons is 10. The fitted manifold and covariance can be further used to compute other geometric metrics, including the Riemannian metric, precision matrix, and Fisher information. These inferred metrics were compared to the ground truth and quantified as the relative estimation error (the difference between estimation and ground truth divided by the ground truth). Dots and error bars represent the median, first, and third quantiles from 10 samplings of the dataset (see Methods). Similar analysis on a 2D synthetic manifold can be found in SI Figure 3.

One straightforward approach to evaluate the quality of spatial coding is by decoding location information from neural states. Good decoding performance indicates good spatial coding. We designed a locally linear classification accuracy to evaluate the quality of spatial coding, formally called as spatial coding accuracy (SCA) (Figure 1B, see Methods). Specifically, at each speed bin value, several locations were randomly sampled. For each sampled location, we created two conjugate boxes with centers near the sampled location but in opposite directions. Data within these two boxes were collected, relabeled as class 1 and class 2, and then split into training and test sets. Next, a logistic regressor was trained to classify the data and was evaluated on the test set. The classification accuracy averaged over all randomly sampled spatial locations is the SCA. SCA measures how well two nearby spatial locations can be distinguished by their corresponding neural states.

For each sampled dataset 𝒟_*s*_, we obtained the SCA values at different speed bins using the method described above. We then developed a Bayesian Linear Ensemble Averaging (BLEA) method to assemble results from different 𝒟_*s*_. Specifically, BLEA first uses Bayesian linear regression to fit metric-speed (metric can be SCA) relation for each 𝒟_*s*_, and then assemble results from different 𝒟_*s*_ by Bayesian averaging^37,38^. The overall BLEA provides a linear relation between metric to speed, taking into account different 𝒟_*s*_. Furthermore, this linear relation is described as distributions (not point estimates), which allows us to compute the confidence interval (CI), p-values and other statistics from a Bayesian framework (see Methods)^39^.

Applying BLEA to SCA (Figure 1C, D), we found that SCA increases with increasing speed, with slope significantly larger than that of label-shuffled dataset. This indicates that grid cell population code improves with increasing speed.

### Gaussian Process with Kernel Regression (GKR) method for fitting manifold and covariance matrices from noisy neural states

What are the underlying neural mechanisms contributing to the improved spatial coding in grid cells? To explore this question, we need a tool to analyze the hard-to-interpret high-dimensional noisy neural states. A recent popular neural population geometry framework suggests that, instead of analyzing noisy high-dimensional data, it will be more intuitive to use certain methods to extract data’s underlying smooth manifold along with noise covariance^34,40,41^. Wishart process is such a method that can infer smooth manifold and covariance matrix^34^. However, the recent implementation of Wishart process requires trial-based experimental paradigm, which forbids this method to be used in broader and complex natural behaving experiments^34^. The OF task (Figure 1A) is one of such natural behaving experiments without strict repeated trials.

Therefore, we developed a novel Gaussian process with Kernel Regression (GKR) method. A dataset (e.g. 𝒟_*s*_) contains noisy neural states ***r*** whose dimensionality equals the number of neurons; and labels ***x*** whose dimensionality equals the number of label variables. A label variable is defined broadly, can be stimulus parameters (e.g. grating stimulus’ orientation), latent variables (e.g. internal decision factor) or behavior variables (e.g.*x, y* locations and speed). GKR assumes that ***r*** follows a Gaussian distribution

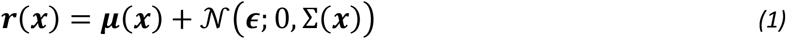

where ***μ***(***x***) is assumed as a smooth-varying mean, also called a manifold in this paper; Σ(***x***) is a smooth-varying covariance. The combination of manifold and noise covariance provides the statistical manifold of neural response, which is a key object in the study in information geometry^27^. The goal of GKR is to infer the manifold and covariance from datasets {***r, x***}.

GKR solves this inference problem by two steps (Figure 2A). In step one, Gaussian process regression is used to infer a smooth ***μ***(***x***) from the data^42^. In step two, residues ***r***(***x***_*i*_) − ***μ***(***x***_*i*_) are computed (index *i* represents *i*th data point), which can be further used to estimate the covariance matrix by kernel averaging (see Methods). Kernel parameters are optimized to maximize data log-likelihood.

### GKR outperforms empirical estimation methods on synthetic datasets

We evaluated GKR on both a one-dimensional synthetic model (Figure 2B, C, D) and a two-dimensional synthetic model (SI Figure 3). Each synthetic model comprises a ground truth manifold ***μ***(***x***), where each component represents a synthetic neural tuning curve; and a covariance matrix Σ(***x***). The synthetic models generate data in accordance with a Gaussian distribution (Equation 1).

**Figure 3:**
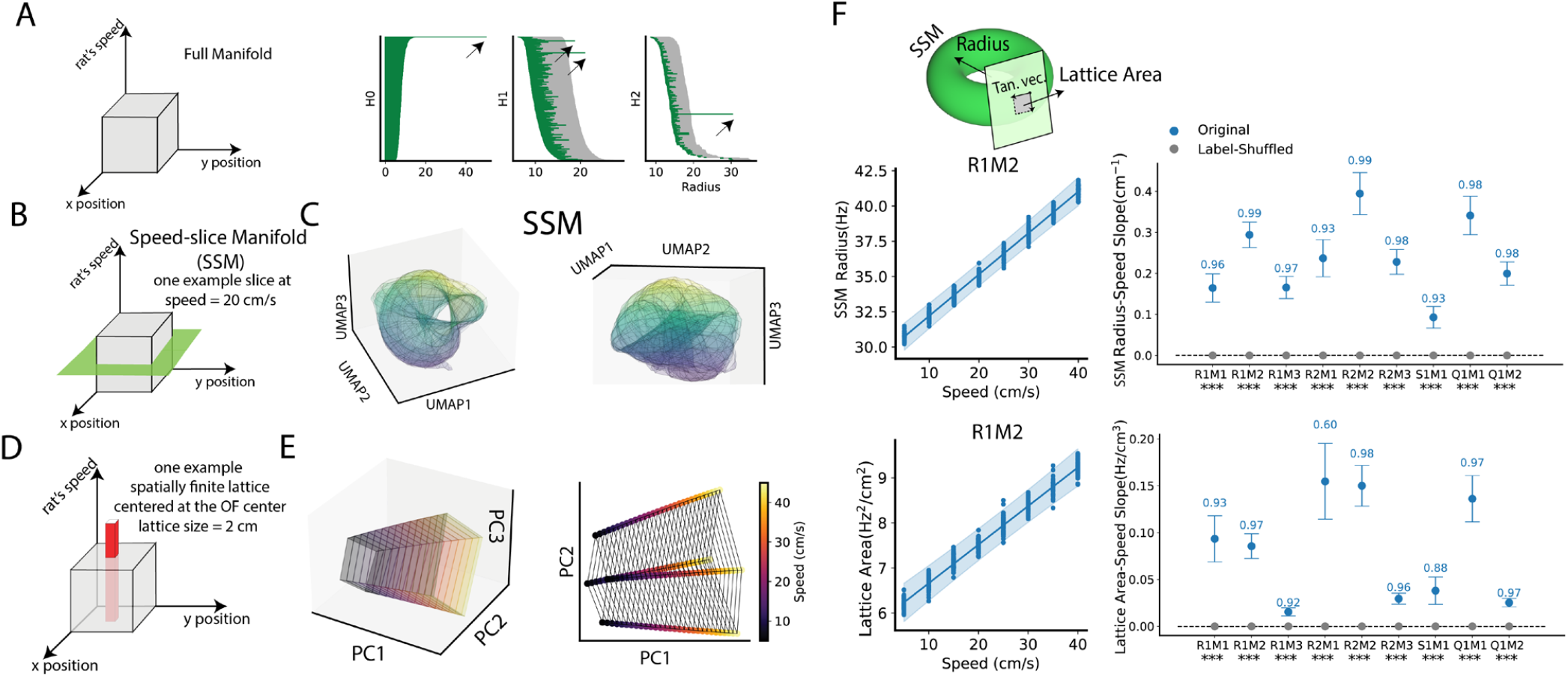
Speed dilates the grid cell population toroidal-like manifold. (**A**) Topological structure of the full manifold. A sampled dataset 𝒟_*s*_ (R1M2) was fed to GKR to fit the manifold and covariance. The resulting manifold is intrinsically three-dimensional (labeled by x, y locations, and speed). Random manifold points were sampled and projected to the first six principal components (PCs) subspace using PCA. Dimensionally reduced manifold points were then subjected to the persistent homology analysis (same analysis but without PCA can be found in SI Figure 4B). In persistent homology, each sampled point on the manifold is surrounded by a small ball with a certain radius (x axis). As the radius increases, some topological structures (holes) emerge and some disappear, as shown by the starting and ending of horizontal bars in the panel. H0, H1, and H2 represent 0D (a whole manifold), 1D (a circular hole), and 2D (a cavity hole) respectively. Bar lengths longer than the grey threshold (maximum bar length in 20 shuffles, see Methods) are considered as long bars, indicated by black arrows. A long bar suggests the existence of a true hole structure in the manifold. A torus has one 0D hole, two 1D holes, and one 2D hole. (**B**) An example speed-slice manifold (SSM) where speed is fixed at 20 cm/s. (**C**) For visualization, the SSM was first projected to the first 6 PCs, then further projected to 3 latent dimensions using UMAP^44^. Color represents the third UMAP component value only for better visualization. The left and right panels show two views of the same SSM. (**D**) We also visualized the representation of four fixed spatial positions (i.e., lattice) but varying speed values. (**E**) For visualization, the lattice manifold was projected into the first three PCs (left) and two PCs (right) respectively (see SI Figure 5 for accumulative variance explained ratio). (**F**) SSM size was measured in the original high-dimensional space (dimension equals the number of neurons). SSM radius is the average distance from points on the manifold to the manifold center. The lattice area measures the parallelogram area formed by two tangent vectors (i.e. Tan. vec., differentiated along x and y labels respectively, see Methods). Left: Each dot is the measured quantity from one GKR fitted from one 𝒟_*s*_ at a speed value. Line and error band show the best linear fitting line and 95% CI using BLEA (see Methods). Right: dots and error bars show the mean and 95% CI of the estimated slopes (using BLEA). The numbers above error bars are r-squared scores. The texts below x axis tick label represents the significance level whether the slope fitted from original data 𝒟_*s*_ differs from that fitted from label-shuffled data (Bayesian method, see Methods); *** p < 0.001; ** p < 0.01; * p < 0.05; NS not significant.

Subsequently, we applied GKR to these generated datasets to infer the ground truth manifold and covariance matrix. In addition to GKR, we used the bin averaging and the Ledoit-Wolf (LW) methods as comparisons. The bin averaging method discretizes the label space ***x*** into small bins, where data within each bin are used to compute the sample mean and sample covariance as the inferred manifold and covariance matrix. The LW method builds upon the bin averaging approach, and applying an additional shrinkage as a regularization technique for better covariance estimation^43^ (see Methods).

Using the inferred manifold and covariance matrix, we can compute other important geometric quantities, including the Riemannian metric, precision matrix, and Fisher information (see Methods). These inferred quantities were compared to the ground truth by evaluating the relative estimation error, defined as the difference between the estimation and the ground truth, divided by the ground truth.

Across various experimental conditions and in both one-dimensional and two-dimensional synthetic datasets, we found that GKR consistently outperforms the bin averaging and LW methods (Figure 2B, C, D, and SI Figure 3).

### Grid cell population activity manifold exhibits a toroidal-like topology

We then applied GKR to the grid cell sampled dataset 𝒟_*s*_ (Figure 1). The inferred manifold is intrinsically three-dimensional due to having three label variables: two locations and one speed. Previous work has shown that grid cell neural states exhibit a toroidal-like topology within the subspace defined by the first six principal components (PCs) in principal component analysis (PCA), where they worked on data clouds^35^. Here, we tested whether this result can be reproduced by our smooth manifold fitted from GKR. We randomly sampled points on the inferred manifold, and then dimensionally reduced them to the first six PCs (more details see Methods). These data points were then subjected to clustering and subsequently fed into persistent homology analysis. Persistent homology analysis replaces data points with small balls of radius *r*. As *r* increases, some topological structures emerge while others die out. The lifetimes are organized as barcodes. A toroidal topological structure should have one 0D hole (i.e., one long bar in H0 space, see Figure 3A and Methods), two 1D holes (e.g. two long bars in H1), and one 2D hole (H2).

Our persistent homology analysis quantitatively reveals that the manifold exhibit toroidal barcode features, indicating that the manifold fitted by GKR has a toroidal-like topology (Figure 3A for R1M2, others in SI Figure 4A). It’s worth noting that this toroidal structure was not confirmed in the original high-dimensional space (SI Figure 4B). Therefore, here we restrict ourselves to use the term “toroidal-like” topology, rather than claiming the manifold is definitively a torus.

**Figure 4.**
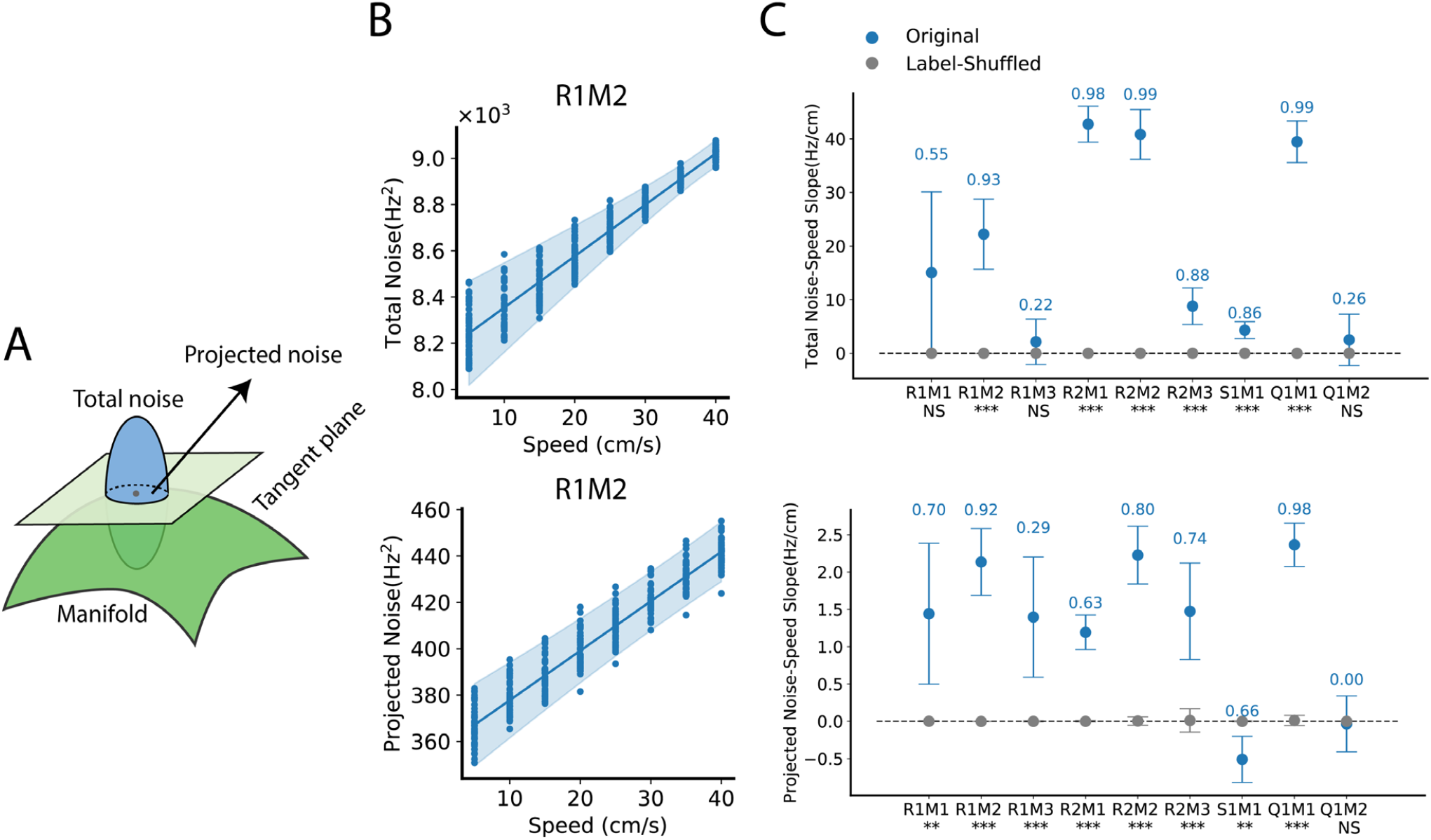
Running speed increases grid cell population noise. **(A)** Total noise is the trace of covariance matrix. Projected noise is the trace of the covariance matrix projected on the SSM tangent plane. (**B**) Left: Each dot is the measured quantity from one GKR fitted from one sampled dataset 𝒟_*s*_ at a speed value. Fifty 𝒟_*s*_ were used. Line and error band show the best linear fitting line and 95% CI using BLEA. (**C**) Dots and error bars show the mean and 95% CI of the estimated slopes (BLEA, see Methods). The texts above error bars represent r-squared scores. The texts below x axis tick label represents the significance level whether the slope fitted from original sampled dataset 𝒟_*s*_ differs from that from label-shuffled data (Bayesian method, see Methods); *** p < 0.001; ** p < 0.01; * p < 0.05; NS not significant.

Given the toroidal-like topology of the full manifold (intrinsically three dimensional), we proceed to examine how it represents spatial locations. For each speed value, we define a speed-slice manifold (SSM, Figure 3B), which is essentially a slice of the full manifold, obtained by holding the speed constant while varying location values. An SSM represents spatial locations information at a fixed speed. For visualize SSM, we randomly sampled points on an example SSM (speed is fixed at 20 cm/s). We then projected these manifold points into the first 6 PC subspace using PCA, and then further projected them into three latent dimensions using Uniform Manifold Approximation and Projection (UMAP)^44^. The resulting visualization (Figure 3C) and persistent homology analysis (SI Figure 4C) suggest this SSM has a toroidal-like structure, as expected from previous analysis of the full manifold (Figure 3A).

### Running speed dilates grid cell toroidal-like speed-slice manifold

A natural question is how the speed modulates the SSM geometry. Visualizing SSMs at different speeds is challenging, so we visualized the speed modulation of an example lattice on the SSM instead. A lattice consists of four spatially nearby points, denoted as ***μ***(***x***_*i*_) where *i* = 0, 1, 2, 3, The spatial components of *x*_*i*_ are at the four corners of a small square in space (centered at the OF center, with a square length of 2 cm), while the speed component of ***x***_*i*_ varies from 5 cm/s to 45 cm/s. PCA applied to the lattice manifold suggests that the lattice manifold is low-dimensional: 3 PCs explain more than 90 percent of the variance (SI Figure 5). Therefore, the lattice manifold was directly projected to three/two dimensions for visualization (Figure 3D, E). It can be observed that the lattice expands with increasing speed. We also visualized other manifold slices, including fixing the rat’s x location, and a larger lattice. All manifold visualizations suggest that the SSM dilates with increasing speed (SI Figure 5).

**Figure 5:**
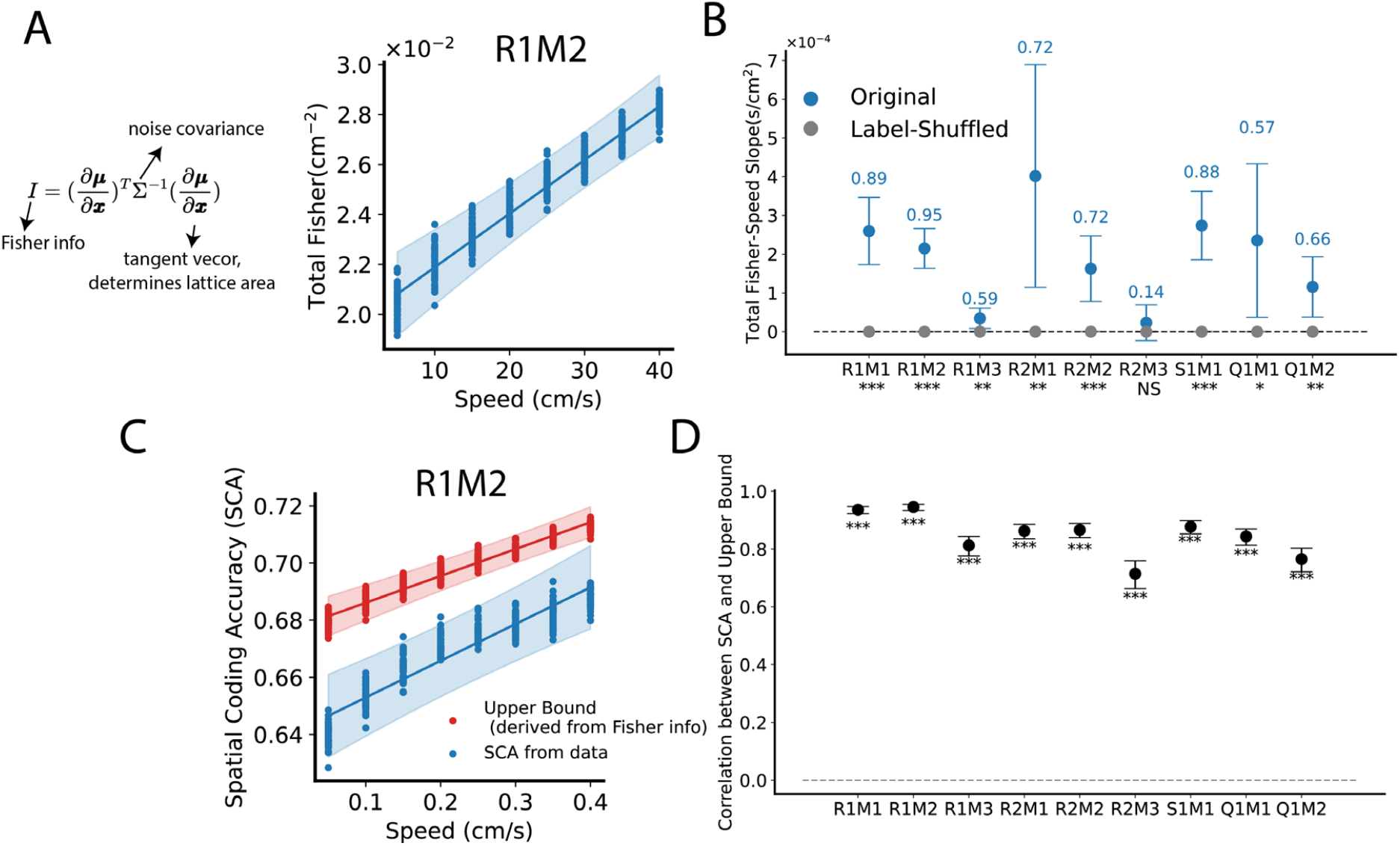
Grid cells’ Fisher information increases with increasing running speed. (**A**) Fisher information mathematically combines noise covariance and tangent vectors (which determine lattice area). It is commonly used to measure the local discriminability of information from noisy neural states^45^. Total Fisher information is the trace of the Fisher information matrix. Each dot represents the measured quantity from one dimensionally reduced sampled dataset 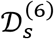 (𝒟_*s*_ projected into its first six PC subspace) at a specific speed value. Fifty 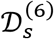 were used. The line and error band show the best linear fit and 95% CI using BLEA. (**B**) Dots and error bars show the mean and 95% CI of the estimated slopes using BLEA (see Methods). The numbers above the error bars represent r-squared scores. The texts below each x-axis tick label indicates the significance level of whether the slope fitted from the sampled dataset 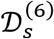 differs from that of label-shuffled data (Bayesian method, see Methods); *** p < 0.001; ** p < 0.01; * p < 0.05; NS not significant. (**C**) We computed theoretical SCA upper bounds based of the Fisher information (see Methods). We also showed the actual SCA computed directly from data (the same as Figure 1B, C). Dots and error bands have the same meaning as in panel (A). (**D**) Correlation between upper bounds and SCA. Dots and error bars represent the mean and 95% CI of the estimated correlation (see Methods). Texts below the error bars indicate the significance levels of whether the correlation differs from zero: *** p < 0.001; ** p < 0.01; * p < 0.05; NS not significance (see Methods). Fisher information is known hard to estimate in a high-dimensional space^24^, therefore all panels in this Figure use the dimensionally reduced dataset 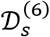. Results obtained by identical analysis on the original datasets 𝒟_s_ are similar, except that, four out of nine experimental conditions do not show statistically significant Fisher information-speed slopes, which may be due to the curse of dimensionality (SI Figure 6).

We quantified SSM size using two metrics. (1) SSM radius is the average distance from the SSM surface to the SSM center. This measures global SSM size. (2) Lattice area is the area of a parallelogram whose two sides are tangent vectors of the SSM. This measures local manifold surface size. Results show that, across all experimental configurations inspected, SSM radius and lattice area consistently increase with increasing speed, indicating that increasing running speed dilates the grid cell toroidal-like SSM (Figure 3G).

### Running speed increases grid cell population noise

The SSM dilation intuitively benefits spatial coding. Consider a binary classification of two classes of neural states representing two nearby locations (e.g. Figure 1B). SSM dilation increases the distance between these two classes, making the representations more distinguishable. However, “distance” is not the only factor determining the quality of spatial coding; another important factor is noise strength. Increasing grid cell population noise reduces discriminability. In the context of spatial coding, this then raises the question to what extent running speed modulates grid cell population noise?

Each sampled dataset 𝒟_*s*_ was used to fit one GKR and to obtain the covariance matrix Σ(***x***). Total noise is the trace of the covariance matrix (Figure 4A). We found that total noise increases with increasing speed (Figure 4B, C). Compared to total noise, noise projected onto the manifold may be more relevant to information coding^26^. Therefore, we projected the covariance matrix onto the tangent plane of the SSM, and the trace of the projected covariance matrix is the projected noise. Consistent with total noise, we found that projected noise also increases with increasing speed (Figure 4B, C).

### Fisher information increases with increasing speed, indicating that the effect of speed-slice manifold dilation outpaces the effect of increasing noise

On the one hand, increased running speed dilates the SSM, pushing neural representations of nearby locations further apart, which benefit spatial coding; on the other hand, increased running speed also raises grid cell population noise, which impairs spatial coding. How does the overall quality of spatial coding change? To address this question, we considered the (Linear) Fisher information. Fisher information is (∂***μ***/ ∂***x***)^*T*^Σ^−1^(∂***μ***/ ∂***x***), which takes into account both the noise factor (Σ) and the lattice area factor (lattice area is formed by tangent vectors ∂***μ***/ ∂***x***) (Figure 5A). Fisher information is commonly used as a metric evaluating the local discriminability of a neural population representation. Total Fisher information is the trace of the Fisher information. Larger total Fisher information implies better spatial coding^45^.

It is well known that Fisher information is hard to estimate in a high-dimensional space^24^. Therefore, besides using the original 𝒟_*s*_, we also projected 𝒟_*s*_ into the first six PCs, denoted as 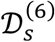. Each 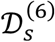 was then fed into GKR for fitting a GKR model. Both GKR fitted 𝒟_*s*_ and 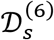 were analyzed identically to double-check our results on Fisher information.

We computed the total Fisher information from the fitted GKRs (see Methods, SI Figure 6A for 𝒟_*s*_ and Figure 5A for 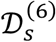). Slope analyses shows that, in both 𝒟_*s*_ and 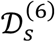, Fisher information increases with increasing running speed (Figure 5B for 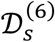, SI Figure 6B for 𝒟_*s*_, although four out of nine experimental configurations in the results of 𝒟_*s*_ do no show statistical significance, possibly due to the curse of dimensionality). The increase in Fisher information with rising running speed suggests that the effect of SSM dilation outpaces that of increasing noise, resulting in improved spatial coding at higher speeds. In fact, Fisher information computed from a purely bottom-up geometric approach is intrinsically related to the SCA (Figure 1B) computed from a top-down decoding approach (see Figure 1B). Specially, we derived the theoretical upper bound of SCA directly from the Fisher information (see Methods). This upper bound is approximately a linear function of the square root of total Fisher information. Hence the increase of Fisher information explains the increase of SCA. We firstly tested this theoretical upper bound in synthetical datasets (SI Figure 7). We show that the SCA computed directly from the synthetical datasets is well bounded by the theoretical upper bound predicted by the Fisher information. Moreover, the trending of upper bounds is consistent with the trending of actual SCA, when varying different dataset parameters (e.g. number of data points, dimensionality etc.)

**Figure 6.**
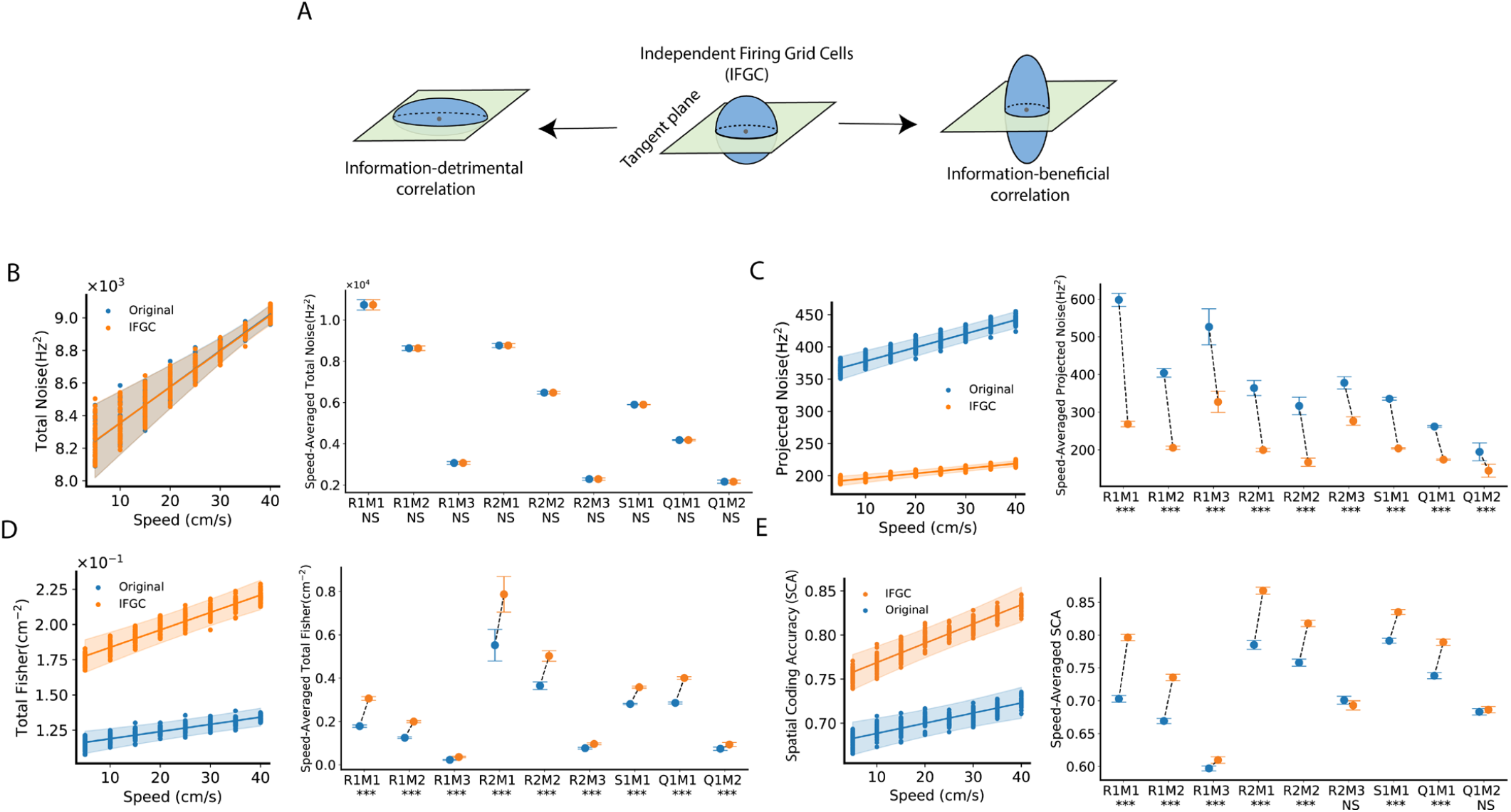
Grid cell noise correlation causes more noise to be projected onto the manifold surface and is detrimental to information coding. (**A**) The presence of noise correlation can result in either better (information-beneficial) or worse (information-detrimental) coding compared to hypothetical independent firing grid cells (IFGC). (**B**) IFGC’s GKR is identical to the GKR fitted from the original sampled dataset 𝒟_*s*_, except that the non-diagonal elements of the covariance matrix are set to zero (see texts and Methods). Total noise was then computed. Left: Each dot represents the total noise from one GKR fitted to one 𝒟_*s*_ at a specific speed value. Fifty 𝒟_*s*_ were used. The lines and error bands show the best linear fitting and 95% CI using BLEA. Right: Speed-averaged total noise is defined as the average total noise across all speeds (from 5 cm/s to 45 cm/s). Dots and error bars show the mean and 95% CI of the estimated speed-averaged total noise (see Methods). The texts below each x-axis tick label indicate the significance level of whether the speed-averaged total noise fitted from the original GKR differs from that of IFGC GKR (Bayesian method, see Methods). *** p < 0.001;** p < 0.01; * p < 0.05; NS not significant. (**C, D**) Same as (B), but measuring projected noise and total Fisher. (**E**) The key idea of SCA is to compute the classification accuracy of data points from two boxes. To compute IFGC’s SCA, we generated a ‘trial-shuffled’ dataset by permuting each cell’s firing across all data points within the same box. This method preserves single-cell firing statistics while disrupting cell-to-cell correlations. Left: We computed SCA using both the “trial-shuffled” data and the original data. Right: Same as the right panel of (BCD).

We further tested the upper bound in grid cell datasets. We calculated the upper bounds of SCA using Fisher information fitted from GKR. The resulting upper bounds are well above SCA (Figure 5C, SI Figure 6C). More importantly, both SCA and the upper bound demonstrate a similar speed modulation effect. The correlations between the upper bounds and actual SCA are statistically significant and positive (Figure 5D, and SI Figure 6D). Overall, Fisher information derived from the geometric properties of the SSM and noise can quantitatively match results from directly measuring decoding performance using SCA. Both approaches support that grid cell spatial coding improves with increasing running speed.

### Grid cell activity noise correlation is information-detrimental

Our results suggest that grid cell spatial coding improves at high-speed, based on our analysis of simultaneously recorded grid cell population activities. One advantage of analyzing simultaneously recorded grid cell activity, compared to individual neural analysis, is that grid cell population analysis implicitly includes the effects of noise correlation on spatial coding. Here we use the term “noise correlation” specifically as the cell-to-cell noise covariance (two different cells). Noise correlation can be information-beneficial or information-detrimental, depending on the geometric relation between noise covariance and information encoding manifold (Figure 6A)^26,46^. In this section, we expose more explicitly the effects of activity correlations on grid cell’s spatial coding.

To explore the role of noise correlation in spatial coding, our main idea is to compare the results from the original grid cell sampled dataset (𝒟_*s*_) to that of hypothetical independent firing grid cells (IFGC)^26^. A classical method of “removing” noise correlation is by trial shuffling. Consider neural states collected under the same experimental condition (e.g., same location), but different trials. To remove noise correlation, one can randomly permute each neuron’s firing profile across trials. This permutation procedure does not change single-cell statistics but “disrupt” the cell-to-cell noise correlation.

Although the OF task has no repeated trials, GKR acts as a generative model, enabling us to generate multiple data points for each condition. This allows us to adapt the “trial-shuffling” approach to obtain IFGC’s GKR. More specifically, given a condition ***x***, the fitted GKR “generates” an infinite amount of data points via a Gaussian distribution (Equation 1). If we apply the permutation procedure above and compute the manifold and covariance again, the resulting mean remains identical, but the covariance will have only diagonal elements, with non-diagonal elements set to zeros. Thus, an IFGC’s GKR model is essentially the original GKR but with purely diagonal covariance matrix.

We computed the total noises from the GKR and IFGC GKR models, respectively, and found that their total noises are identical (Figure 6B). This result is expected, as removing the non-diagonal terms does not alter the trace of the covariance matrix. However, it changes the projected noise. As shown in Figure 6C, the projected noise from the GKR model is larger than that from the IFGC GKR model. This suggests that the cell-to-cell noise correlation in grid cell activity “reshapes/reorients” the noise structure, leading to more noise being projected onto the torus surface. Further comparison of Fisher information revealed that the presence of noise correlation reduces Fisher information, indicating that noise correlation is information-detrimental (Figure 6D).

To double-check this, we used a completely top-down decoding approach. Recall that the key idea of SCA is computing the linear classification accuracy of neural states from two boxes. To obtain IFGC SCA, we randomly permute each neuron’s firing rates within each of the boxes. This does not change single-cell statistics in a box but “disrupt” the noise correlation. We computed IFGC SCA from this permuted box data, showing that IFGC SCA is larger than SCA (Figure 6E). In other words, the presence of noise correlation reduced SCA, confirming that the effect of noise correlation is information-detrimental.

## Discussion

In navigation, it’s important to maintain internal spatial representations while the animal is moving. Grid cell is thought as a fundamental building block in internal spatial representation^2,3^. Prior analyses of speed modulation on grid cell coding have primarily focused on individual cells or cell pairs^11–14^. These approaches, however, did not account for population noise covariance, a factor that can substantially influence coding fidelity^26^. Here, we developed GKR to study the population coding from an information geometry perspective. We demonstrated that grid cell manifold expands in size as speed increases. This manifold dilation effect surpasses increase in noise, as indicated by higher Fisher information observed at high speeds. Overall, our results favor the hypothesis that increasing running speed increases grid cell spatial coding accuracy. GKR can be a powerful tool to study neural population representation from an intuitive information geometric perspective.

Besides grid cells, does running speed enhance the information representation of other cell types within the navigation system? Hardcastle et al found that, the MEC exhibits more neurons encoding spatial information at high speeds (10-50 cm/s) than at low speeds (2–10 cm/s)^10^. Furthermore, the spatial decoding accuracy of MEC neurons shows improvement at higher speeds, indicating that increased speed generally benefits MEC spatial representation^10^. This is supported by the fact that, similar to grid cells, running speed mostly increases the firing rates of other cell types in the MEC (e.g., head direction cells, speed cells, conjunctive cells)^11,17^ and hippocampus (e.g., place cells)^47^. The increase in firing rates can closely relate to the manifold dilation shown in Figure 3, since each component of the manifold represents a single neuron’s tuning^41^. Therefore, the speed dilation phenomenon shown in this paper may also apply to other cell types in the navigation system. Although, more work needs to be done to compute the noise covariance structure to calculate the Fisher information, which is more directly related to spatial coding.

Beyond the navigation system, running speed modulation effects have been widely observed across the brain^48^. For example, locomotion primarily suppresses neural activities in the auditory cortex^49,50^. On the other hand, locomotion generally enhances V1 neuron activity^51,52^, but may turn to suppression after certain high running speeds^53^. In fact, the effect of locomotion modulation is usually entangled with other modulation factors^48,53,54^. For example, V1 neural activities are jointly influenced by both animal’s running speed and visual stimuli movement speed^52^. This influence can be mathematically expressed as a weighted sum of the two speed contributions, with weights varying diversely across neurons.

Geometric approach has been shown to be a practically effective approach to assist understand the diversity of individual neuron from a comprehensive population level^40,55,56^. GKR can be a useful tool to understand the diversity of running speed modulations in different brain areas.

One advantage of GKR is its ability to provide detailed inspection of location geometry. Local geometry reveals the intricacies of information coding within a small range of values, which is particularly useful for comparing the representation bias of different information values. Representation bias has been observed in the navigation system^47,57^. For instance, place and grid cells’ fields tend to shift towards reward locations, which was interpreted as an over representation to rewarded locations^57–60^. From a geometric perspective, over representation implies larger Fisher information, which can be attributed to either local manifold dilation, reduced projected noise, or both (Figure 3, 4, 5). The concepts of local manifold dilation and reduced noise have been supported in working memory studies: (1) The working memory system may use attractors to represent high-probability (high-prior) information values^61–63^. Attractors attract nearby neural states, therefore reduce random diffusion noise, hence benefits Fisher information^64^. (2) Recurrent neural networks (RNNs) trained on working memory tasks utilize larger state spaces to represent high-prior values, thus also benefits Fisher information (larger tangent vector length)^61^. In RNNs, manifolds are often observed to be quite simple, usually taking the form of a low-dimensional ring structure^61^. This simplicity allows the size of the encoding space to be measured using straightforward methods. However, in the actual brain, manifolds can be highly complex and high-dimensional^56^. The GKR method illustrated in this paper can be particularly helpful in studying the local structure of these complex, high-dimensional manifolds, assisting the analysis of representation bias.

Neural activities are known to exhibit correlations. The presence of noise correlation can be either information-beneficial or information-detrimental, depending on the fine structure of noise covariance and information encoding geometry^26^. Inferring noise covariance is challenging. It is important to note that many studies utilize artificial, simplified, and low-dimensional stimuli, which allow for precise replication of experimental conditions and trial repetition^34,46^. In contrast, real-world information is typically high-dimensional, less controllable, and lacks repeated trials^31,33^. This presents a significant challenge in computing real-world information coding. One approach to address this challenge involves modeling neural systems using explicit neural network models and then computing information coding from these models^36^. While this method has shown promise in retinal systems, even the most advanced deep neural network models struggle to fully capture the complexities of biological systems^65^. GKR offers an alternative, neural-network-model-free approach to study information coding in naturalistic stimuli and behavior. However, GKR is not omnipotent. It will still face limitations when dealing with extremely high-dimensional data. This challenge can be mitigated by preprocessing data using dimension reduction methods^66^. In general, the combination of dimension reduction techniques with manifold inference methods (such as GKR and Wishart processes^34^), presents a promising avenue for exploring neural coding in complex, high-dimensional, and naturalistic information contexts.

## Methods

### Experimental Data

Experimental data were collected by Gardner et al.^35^. Rats performed open-field foraging (OF) tasks in a 150 cm wide OF box. Three-dimensional motion capture tracked the rats’ head positions and orientations using five retroreflective markers attached to the implant during recordings. The 3D marker positions were then projected onto the horizontal plane to determine the rats’ 2D positions. Neuropixel probes recorded neural activity in the MEC. Neural activities were then processed using a clustering method to classify neurons into grid cells and non-grid cells^35^. In total, these procedures yielded nine sets of simultaneously recorded grid cell population activities: rat ‘R’ day 1 modules 1, 2, 3; rat ‘R’ day 2 modules 1, 2, 3; rat ‘S’ module 1; and rat ‘Q’ modules 1, 2. These are also called data under nine experimental configurations in this paper. We used a shorthand notation, e.g. “R1M2”, to represent rat R (“R”) on day 1 (“1”) and grid cell module two (“M2”). Note “R1” does not necessarily means the same day as “S1”. Day labels are only to distinguish the same rats. These processed data are available from Gardner et al. 2022^35^.

### Grid Cell Rate Map

The grid cell rate maps shown in Figure 1A and SI Figure 1 were computed as follows. Firing rate was estimated by spike counts divided by 10 ms time bins and then temporally convolved with a Gaussian filter of (standard deviation Σ = 20 ms). To estimate the averaged firing rate at different locations, the OF box (150 × 150 cm) was digitized into 3 × 3 cm small spatial bins. Firing rates at each visited spatial bin were averaged, and those at each unvisited bin were set to 0. To correct the effect of unvisited bins, we created a mask (M_0_) with a value of 1 at the visited bins and 0 at unvisited bins. Next, both the firing rate and mask M_0_ were spatially convolved with a 2D Gaussian filter of Σ = 8.25 cm. The convolved firing rate was divided by the convolved M_0_ to obtain the final corrected rate map for each cell.

### Gridness

Gridness measures how well a grid cell’s rate map conforms to a hexagonal pattern^12^. Some grid cells’ rate maps have incomplete peaks at the OF box boundaries. To correct this boundary effect, the rate map was first padded by 30 cm on each side. This padding was done by linearly ramping the edge’s firing rate map to zero at the farthest 30 cm padded position (implemented using the ‘numpy.pad’ function in Python, with ‘mode=‘linear_ramp’’). Autocorrelating the padded rate map produced an autocorrelation map. The boundary effect of the autocorrelation was corrected by padding zeros on all sides (implemented using ‘scipy.signal.correlate2d (padded_rate_map, padded_rate_map, mode=‘same’, boundary=‘fill’, fillvalue=0)’).

The autocorrelation map was masked by two circles centered at the map’s center. The outer circle’s diameter matched the edge length of the autocorrelation map. The inner circle’s area was 15% of the outer circle’s area to filter out center peaks on the map. Only the regions between the two circles were kept; the rest were set to 0. Next, the masked autocorrelation map was correlated with itself after rotating 30, 60, 90, 120, and 150 degrees, respectively. A well-defined grid cell should have peak correlation values at 60 and 120 degrees, and valleys at 30, 90, and 150 degrees. Gridness was calculated by subtracting the average valley values (30, 90, and 150 degrees) from the average peak values (60 and 120 degrees).

### Data Preprocessing

Time was binned by every 10 ms. Spikes count at each time bin was computed, and then divided by 10 ms as an estimation of firing rate. The firing rate was then temporally smoothed using a Gaussian kernel with a standard deviation of 20 ms. To estimate the rat’s speed, velocity was firstly computed as the finite differences of rat positions, i.e. (***P***_***i***+**11**_ − ***P***_***i***−**11**_)/20 where ***P***_***i***_ is the rat’s position at time bin *i*. The velocity’s L2 norm is the speed. This procedure provided a feature map indicating grid cell firing rates, with rows representing time bins and columns representing grid cell IDs; and a label with rows representing time bins and three columns indicating *x* location, *y* location, and speed. Data (feature map and label) with too low (speed < 5 cm/s) or too high (> 45 cm/s) speeds were excluded. Grid cells with low gridness (below 0.1) were also excluded. The final feature map and label combined are termed a grid cell dataset, denoted as 𝒟. There are 9 grid cell datasets corresponding to different experimental conditions (different rats, grid cell modules, and different days). The number of grid cells in each dataset is: 113 in R1M1, 132 in R1M2, 51 in R1M3, 140 in R2M1, 153 in R2M2, 62 in R2M3, 96 in S1M1, 81 in Q1M1, 53 in Q1M2.

The speed distribution in 𝒟 is highly biased. It has more data in low-speed region than in the high-speed region (SI Figure 2). This biased distribution of data may cause potentially biased evaluation. To avoid this, we performed resampling on the dataset as follows. Speed ranging from 5 cm/s to 45 cm/s was binned into 5 cm/s bins. Data in each speed bins were collected. Denoting the minimum number of data points among all speed bins as 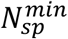. Then we defined *K* as the 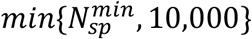. In each speed bin, we sampled *K* data points (without replacement). Sampled data points from different speed bins were combined to create a single sampled dataset, denoted as 𝒟_*s*_. 𝒟_*s*_ has approximately fair amount of data at each speed value. The above sampling procedure was repeated 50 times, resulting 50 sampled datasets 𝒟_*s*_.

As a baseline comparison, we also shuffled the data 𝒟 by permuting the label timestamps, thereby disrupting the relationship between neural states and labels. This permuted data was then processed using the same sampling procedure as described above, yielding 50 label-shuffled-sampled datasets.

The dimensionality of 𝒟_*s*_ is the number of grid cell, which can be more than 100. This can bring challenge in accurately estimate covariance and Fisher information^24^. Therefore, we also performed PCA on 𝒟_*s*_, projecting to the first 6 principal components to obtain 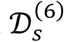. Same projection procedure was also applied in the shuffled datasets. Throughout this paper, these projected datasets were only used for estimating Fisher information and comparing SCA (i.e. Figure 5).

### Spatial coding accuracy

A common way to evaluate the quality of neural population representation is to assess how accurately a simple linear classifier can distinguish between neural population representations of two adjacent experimental conditions (e.g., stimulus parameters or locations in this paper)^22^. In this paper, this type of classification accuracy is referred to as spatial coding accuracy (SCA, Figure 1B).

Consider one sampled dataset 𝒟_*s*_, we split it into 8 speed-split datasets (SSD) based on speed values. Specifically, data with speed values within [*v*_*i*_, *v*_*i*_ + 5cm/s] were collected as one SSD, where *v*_*i*_ = 5,10, …, 40 cm/s. For each SSD, we randomly sampled 300 spatial locations ***x***_*c*_. For each location ***x***_*c*_, we constructed two adjacent locations ***x***_±_ = ***x***_*c*_ ± *δl* ê, where ê is a unit vector with a random angle, *δll* = 5 cm. Each ***x***_±_ defines a small spatial box, centered at ***x***_±_ with an edge length of 10 cm. Data within two boxes were collected. To ensure fair classification, the dataset with the larger number of data points was subsampled (without replacement) so that both boxes had an equal number of data points. Data from two boxes were then concatenated. If the total number of data points was less than 50, this ***x***_*c*_ was discarded due to insufficient data. Otherwise, the concatenated data was split into train and test sets (0.67:0.33). A logistic classifier (with a L2 regularization coefficient C = 1, implemented by the scikit-learn package) was then trained on the train set and evaluated on the test set. The classification accuracies averaged across all valid ***x***_*c*_ is the SCA of that speed bin [*v*_*i*_, *v*_*i*_ + 5cm/s]. This procedure was applied to all speed bins, 𝒟_*s*_ (or 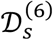in Figure 5C), and label-shuffled dataset.

### Bayesian linear ensemble averaging and statistical testing

Metrics considered in this paper include SCA (e.g. Figure 1C, D, 5C), torus radius, lattice area (e.g. Figure 3F), total and projected noise (e.g. Figure 4B, C), and Fisher information (e.g. SI Figure 6A, B), etc. For each sampled dataset 𝒟_*s*_, we computed the metric values at different speed bins, forming a metric-speed dataset consisting of metric value *t*_*i*_ and the corresponding speed value *v*_*i*_, where *i* indexes the *i*th data points in the metric-speed dataset. For example, one dot in Figure 1C is one data point in the SCA-speed dataset (with a corresponding 𝒟_*s*_). For convenience, we also wrote ***x***_***i***_ = (*v*_*i*_, 1), which includes speed and a constant for a bias parameter. Currently we limit our discussion to one 𝒟_*s*_, and later we will ensemble results from different 𝒟_*s*_ by Bayesian model averaging^38^.

Given a metric-speed dataset from one 𝒟_*s*_, we used Bayesian linear regression (BLR) to fit the linear relationship between a metric and speed^42^. The benefit of BLR over Ordinary least squares is that BLR naturally provides a way to set the regularization parameter (by setting the prior) and offers the posterior distribution of inferred parameters (e.g., slope), allowing analyzing the data from a pure Bayesian perspective. We follow the implementation of BLR in Bishop 2006^42^.

In BLR, the relationship between metric and speed is modeled as

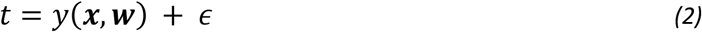

Where ϵ ∼ 𝒩(ϵ| 0, *β*^−1^), *β* is a scalar representing precision, and *y* = ***w***^*T*^***x***. This equation equivalently tells the conditional distribution *p*(*t*|***x, w***) is a Gaussian distribution with mean *y* and variance *β*^−1^. The prior of parameter ***w*** is modeled as ***w*** ∼ 𝒩(***w***|0, *α*^−1^𝕀) where 𝕀 is a 2 × 2 identity matrix, and *α* is a scalar. Given the prior and conditional distribution, we can derive the posterior distribution *p*(***w***|***t***, *X*) and predictive distribution *pt*_*q*_***x***_*q*_, ***t***, *X* where ***x***_*q*_ is the query label, ***t*** and *X* are the data points in the metric-speed dataset, *t*_*q*_ is the prediction. Both posterior and predictive distribution are Gaussian distributions. Hyperparameters *α* and *β* were estimated by maximzing the marginal likelihood *p*(***t***|*α, β, X*) through an iterative method^42^. Overall, BLR provides the posterior distribution *p*(***w***|***t***, *X*) and predictive distribution *pt*_*q*_***x***_*q*_, ***t***, *X* from one metric-speed dataset (obtained from one sampled dataset 𝒟_*s*_). To simplify the notation, the two distributions are written as *p*(***w***|𝒟_*s*_) and p*t*_*q*_***x***_*q*_, 𝒟_*s*_), which follows 𝒩***w***| ***m***_*w,s*_, Σ_*w,s*_ and 𝒩*t*_*q*_| *m*_*t,s*_, Σ_*t,s*_, respectively.

We ensemble results from sampled datasets 𝒟_*s*_, i.e. *p*(***w***|𝒟) and p averaging^38^. Since each 𝒟_*s*_ is a random subsampling (under a “equal amount of data points in each speed bin” restriction, see Methods: Data Preprocessing) of the 𝒟, *p*(***w***|𝒟) = ∑_*s*_ *p*(***w***|𝒟_*s*_)*p*(𝒟_*s*_ |𝒟) = ∑_*s*_ *p*(***w***|𝒟_*s*_)/ *B* where *B* = 50 is the total number of sampled datasets. *p*(***w***|𝒟) is a mixture of Gaussian distributions, for simplicity, we approximated it into a single Gaussian distribution with the same mean and covariance (see details in SI Methods). The mean of *p*(***w***|𝒟) is

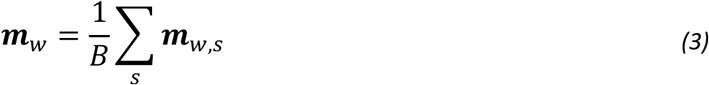

The covariance is

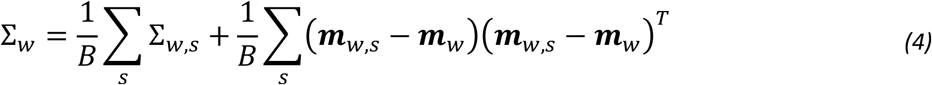

where the first term is the average of covariances, second term represents to bias error. Similarly, p(*t*_*q*_|***x***_*q*_, 𝒟) can be approximated by a Gaussian distribution with certain mean and covariance matrix (see SI Methods). In fact, the mean and covariance matrix are in forms that, after this Gaussian approximation on 𝒟, *t*_*q*_ is still a linear function of ***w***, as can be explicitly written as below

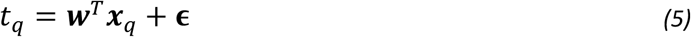

where ***w*** ∼ 𝒩(***w***| ***m***_*w*_, Σ_*w*_) and 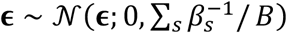 where *β*_*s*_ is the best hyperparameter fitted using iteration method in a sampled dataset 𝒟_*s*_ (see above). Overall, we obtained *p*(***w***|𝒟) and p*t*_*q*_|***x***_*q*_, 𝒟), which are both approximately Gaussian distributions. This overall method pipeline is called Bayesian linear ensemble averaging (BLEA) in this paper. Mathematical details can be found in SI Methods.

*p*(***w***|𝒟) and p(*t*_*q*_|***x***_*q*_, 𝒟) allow us to estimate confidence interval and state statistical significance from Bayesian framework^39^. First, since the predictive distribution (p*t*_*q*_|***x***_*q*_, 𝒟) is a Gaussian, the 95% confidence interval (CI) of the prediction (two-tailed) is given by an interval [a, b] such that Φ(*a* − *μ*) / *σ* = 0.025 and Φ(*b* − *μ*)/*σ* = 0.975, where Φ(·) is a cumulative density function of a standard Gaussian distribution, *μ* and *σ* are the predictive distribution mean and standard deviation. The visual goodness of CIs in Figures (e.g. Figure 1C) covering data points support the validity of BLEA.

95% CI is also called as creditable interval in Bayesian framework^39^.

Similarly, knowing the posterior distribution of slope *p*(***w***|𝒟), 95% CI can also be computed accordingly.

We are interested in whether the slope fitted from 𝒟 is statistically different from that fitted from the label-shuffled dataset. Therefore, we also prepared label-shuffled 𝒟_*s*_ from 𝒟 (see Methods: Data Preprocessing), and run the above analysis to obtain their label-shuffled posterior and predictive distributions. Since the posterior distributions of the original and label-shuffled datasets are both Gaussian, we defined the slope difference *d* = *w*^*data*^ − *w*^*shuffle*^, which follows also a Gaussian distribution with mean as the difference of two slope means and variance as the sum of two variances. Based on the distribution of *d*, probability of direction, *p*_*d*_, can be computed as the maximum of *P*(*d* > 0) and *P*(*d* < 0). *p*_*d*_ tells the probability of *d* to be positive or negative (depending on which is the most probable). It directly relates to p-value (from a frequentist framework, two-sided) by *p* = 2 × (1 − *p*_*d*_), where the null hypothesis is that *d* = 0 and alternative Statistical statement hence can be made based on the p-values.

We are also interested in whether the speed-averaged metric computed under the 𝒟 is statistically different from that computed under the hypothetical independent firing grid cell assumption (IFGC, Figure 6). For each 𝒟_*s*_, we averaged the metric value across speed values. This gives one 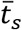. Fifty 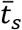 were concatenated and fitted by a Gaussian distribution by maximum log-likelihood, as an approximation of 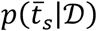 the speed-averaged metric computed under original dataset statistically different from which from the IFGC (Figure 6B, C, D, E).

### Bin Average and Ledoit-Wolf Estimator

We approximated the neural population responses (neural states for short) as a Gaussian distribution:

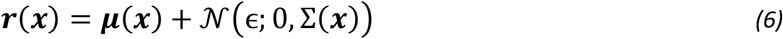

where ***r*** ∈ ℛ^*N*^ represents a neural state contains *N* neurons, and *x* ∈ ℛ^M^ represents *M* labels. Labels are defined broadly. It can be stimulus parameters (e.g., grating image orientation, object positions), an agent’s latent state (e.g., latent dynamics factor, emotion), or an agent’s behavior labels (e.g., agent speed, agent position). ***μ*** is the mean of neural state, modeled as a continuous function of the labels. ***μ*** is also called as a manifold in this paper. ϵ is a white noise with a covariance Σ(***x***). Given noisy neural states ***r*** and corresponding labels ***x***, our goal is to infer the smooth varying manifold ***μ*** and covariance Σ.

Bin averaging is a straightforward estimation method. This approach divides the entire range of label ***x*** into small bins. Data points ***r***_*i*_ within each bin are considered to have an identical label ***x***_*i*_. Hence, the manifold can be estimated by sample average 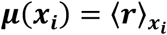, where 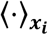 denotes averaging over the data points within bin ***x***_***i***_. Similarly, the covariance Σ can be estimated by sample covariance matrix.

However, when the number of data points in each small bin is sparse and the neural state dimensionality is high (i.e., a large number of recorded neurons), bin averaging can lead to unreliable—and sometimes even non-invertible—estimation of the covariance matrix^24^. To address this, the shrinkage method was proposed. This method is equivalent to adding L2 regularization to the maximum likelihood estimation of the covariance matrix, guiding the estimation towards a more structured assumption (e.g., an identity matrix)^43^. In particular, this paper uses:

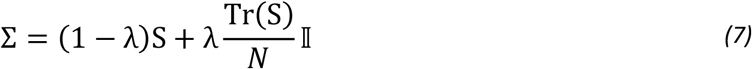

where *S* is the sample covariance, λ is the shrinkage coefficient estimated by the Ledoit-Wolf (LW) shrinkage algorithm^43^, *N* is the number of neurons, and 𝕀 is an identity matrix. This algorithm is implemented by a Python function ‘sklearn.covariance.LedoitWolf’.

### Gaussian Process with Kernel Regression

One disadvantage of the bin average and LW methods is that the estimation of one bin’s covariance does not use data from adjacent bins. Ideally, the manifold and covariance matrix are smooth over label values. Data in adjacent bins can provide certain information about the current bin. Therefore, we developed the Gaussian Process with Kernel Regression (GKR) manifold and covariance from noisy neural states. GKR has two major steps: step 1 is for inferring manifold while step 2 is for inferring covariance matrix.

In step 1, each component of ***r*** across all time bins is standardized to have a mean of zero and a variance of one. Denoting the standardized ***r*** as 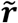. 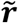then is modeled as 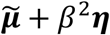, where ***η***is a standard gaussian noise, and *β* is a scalar parameter. The manifold 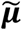 is modeled as an N-independent Gaussian process written as 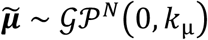 i.e., with zero mean and a kernel function *k*_μ_: ℛ^M^ × ℛ^M^ → ℛ to control the “closeness” of 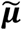 given two different labels ***x***^42^. A shared kernel for all components of 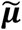 is used in this paper. Although the kernel is shared by all components 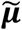, it has different parameters for different components of the label, i.e., 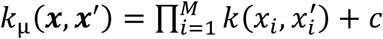 where *x*_*i*_ is the *ii*th component of a label ***x*** and *c* is a constant parameter. The kernel for *x*_*i*_ is

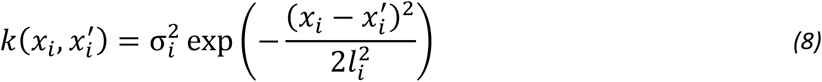

where Σ_*i*_ and *l*_*i*_ are parameters. If *x*_*i*_ is a circular variable, a sine wrapping is applied:

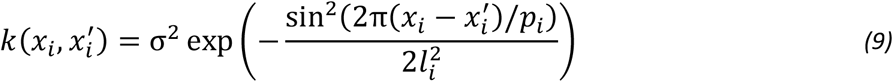

where *p*_*i*_ represents the period of the circular variable. Based on all these modeling, the problem of inferring 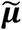 from noisy data 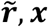 becomes a classical Gaussian process regression problem, where the parameters {*β, l*_*i*_, Σ_*i*_, *c*_*i*_} are optimized to maximize the log-likelihood of a joint Gaussian distribution for 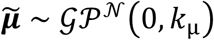 Finally, 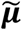 is unstandardized back to ***μ***.

In many scenarios, the label ***x*** spans a large continuous range rather than a few discretized values (e.g., possible positions of a rat in a navigation task). In this case, Gaussian process regression requires computing a large kernel matrix, leading to expensive matrix manipulations^67^. To reduce this, we employed a variational inducing variable method^67^. It approximates training label values with a smaller set of inducing points *z*, thereby reducing the time complexity. In this paper, inducing points were initialized as a randomly sampled subset of the original training labels (200 inducing points), and were optimized during the optimization of Gaussian process regression. Gaussian process regression with inducing variables method is implemented in the Python GPflow package^68^.

The above step one infers the manifold ***μ***(***x***). Step two infers the covariance matrix Σ(***x***). Define the gram matrix of a point (***r***_***i***_, ***x***_***i***_) as *C*(***x***_***i***_) ≡ (***r***_***i***_ − ***μ***(***x***_***i***_)))^***T***^(***r***_***i***_ − ***μ***(***x***_***i***_))) matrix at ***x*** as

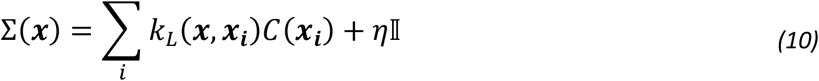

where *i* sums over all training data points, and *η* = 10^−6^ is a small number for numerical stability (keep the covariance invertible even in the first term is small). *k*_*L*_(***x, x***_***i***_) is a weight kernel that represents the contribution of *C*(***x***_***i***_) in estimating the covariance matrix at label ***x***. It is normalized such that ∑_*i*_ *k*_*L*_(***x, x***_***i***_) = 1. To gain an intuition of this method, consider a simple case where (up to a normalization) *k*_*L*_(***x, x***_***i***_) = 1 if ||*x*_*i*_ − *x*|| < *δ* and zero otherwise, step two is simply a sample covariance in a small bin of width *δ*.

Since we assumed covariance is a smooth function over ***x***, *C*(***x***_*i*_) of adjacent ***x***_*i*_ should still contribute to the estimation of Σ(***x***). Therefore, we use a gradually decaying weight kernel

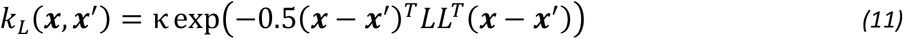

with κ as a normalization Σ_*i*_*k*_*L*_(***x***,***x***_***i***_)=1, and *L* is an M ×M upper triangular matrix, interpreted as the Cholesky decomposition of a semi-positive definite precision matrix *L*^*T*^. Note that the precision matrix has non-diagonal terms, hence the interactions between different label components are considered.

Parameter *L* is optimized to maximize the Gaussian log-likelihood of the data

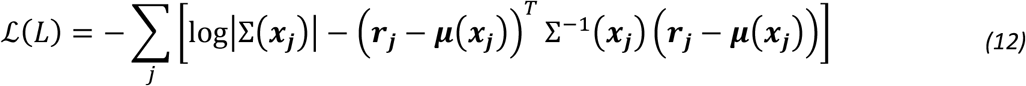

where terms irrelevant to covariance are omitted. Notably, while Σ is the weighted average of the Gram matrices from the training set, the log-likelihood function ℒ should be evaluated from the validation set, where we used different indices *i, j* to distinguish. Setting the log-likelihood function on the training set would result in Σ(***x***) converging to the Gram matrix *C*(***x***). This can be demonstrated by computing Σ(***x***) to satisfy the condition ∂ℒ/ ∂Σ = 0. Therefore, splitting between training for computing covariance and validation for computing likelihood is necessary.

Overall, we use the following procedure to fit the manifold and covariance from a dataset. In step one, the entire dataset was used for Gaussian process regression, obtaining a continuous manifold function ***μ***. In step two, we used batch training. The dataset was split into batches, each containing 3000 data points (except for the final batch). Each batch was further split into train and validation sets (0.66:0.33). The train set was used for computing the covariance matrix given an *L* (initialized as an identity matrix), and the validation set was used to compute the log-likelihood function. The log-likelihood was then maximized by an Adam optimizer (gradient applied on *L*). This batch training was repeated for 30 epochs. Finally, with the optimized *L*, the whole dataset was used for computing covariance (Equation 10).

### One-Dimensional and Two-Dimensional Synthetic Datasets

Synthetic datasets are modeled as Gaussian distributions

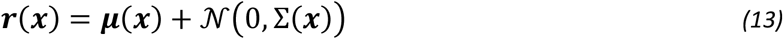

The covariance matrix is Σ(***x***) = *L*^*T*^ where

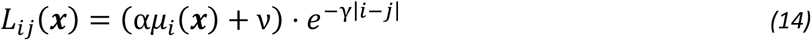

with ν representing a constant for stimuli-independent noise, and *γ* as the non-diagonal decay rate. Note that the covariance matrix Σ depends on the manifold ***μ***.

In the one-dimensional synthetic model, *i*th component of the manifold ***μ*** is given by

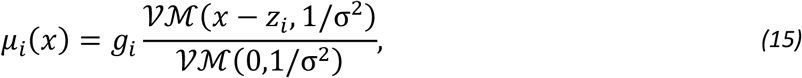

where 𝒱ℳ(·) denotes a von Mises function, *g*_*i*_ ∼ *U*(0.5,1.5) is a random gain, *z*_*i*_ = 2*i*π/*N* is the preferred label value for the *i*-th neuron, and *σ* = 0.3 is the tuning width. *x* is a circular scalar label ranging from 0 to 2π. Parameters for generating covariance matrix (Equation 14) are: *α*= 0.2,*v*= 0.05, *γ* = 1.

In the two-dimensional synthetic model, the *i*th component of manifold ***μ*** is

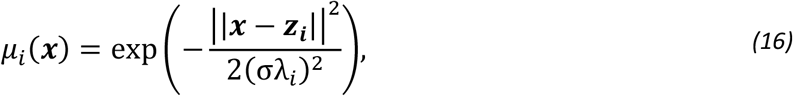

where *z*_*i*_ ∼ *U*([−1,1], [−1,1]) is the center of the receptive field of neuron *i*, Σ = 0.3, and λ_*i*_ ∼ *U*(0.5, 1.5) controls the tuning width. The label ***x*** is two-dimensional, with each component ranging from -1 to 1. Parameters for generating covariance matrix (Equation 14) are: α = 0.5, ν = 0.1, *γ* = 1.

To generate a synthetic dataset of size *T, T* labels ***x*** were uniformly sampled from the entire range. Each sampled label ***x*** was then used to compute one manifold point ***μ*** and one covariance matrix Σ, thus generate one ***r*** using a Gaussian distribution (Equation 13). *T* labels generate *T* data points.

When visualizing the ground truth of synthetic datasets, 100 labels *x* were randomly sampled. Then manifold points *μ* and covariance matrices were computed. Manifold points were then fed into a PCA, dimensionally reduced to the first 2/3 dimensions (two for Figure 2 and three for SI Figure 3). The covariance matrices were also projected onto the PCA subspace, transforming to a 2×2/3×3 matrix. The eigenvalues of this 2×2/3×3 matrix were visualized as the lengths of the ellipsoid’s major axes (Figure 2); and eigenvectors were visualized as the ellipsoid’s major axes directions.

### Computing the relative prediction error of a metric to the ground truth in the synthetic dataset

We evaluated the performances of estimators (Bin average, LW, GKR) by comparing their predictions to ground truth. We evaluated several metrics: (1) manifold ***μ*** (2) covariance matrix Σ (3) Riemannian metric (∂***μ***/ ∂***x***)^*T*^(∂***μ***/ ∂***x***) (4) Linear Fisher information (∂***μ***/ ∂***x***)^*T*^Σ^−1^(∂***μ***/ ∂***x***) (5) Precision matrix Σ^−1^. ∂***μ***/ ∂***x*** was estimated numerically by finite difference.

For each configuration (number of data points or number of neurons, Figure 2), ten synthetic datasets were sampled. For each dataset, all data were used for training the estimator. Trained estimator predicts the values of metrics at other 100 randomly sampled labels. The relative estimation error is the mean of 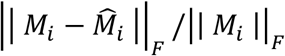 over all 100 label *i*, where M_*i*_ is the ground truth quantity while 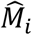 is the estimated quantity, || · ||_*F*_ is the Frobenius norm.

### Fit and visualize grid cell population manifold

GKR was applied to 𝒟_*s*_ to fit manifold and covairance matrix. Since 𝒟_*s*_ has three labels (two for locations and one is the speed), the full manifold is an intrinsically three-dimensional object. For the ease of visualization, we visualized slices of the manifold instead. First, we visualized the manifold representing different locations but fixing the speed at 20 cm/s, i.e. speed-slice manifold (SSM). A 30 by 30 grid of positions was sampled from the entire OF space. Along with the fixed speed of 20 cm/s, these labels were fed into the fitted GKR to predict the manifold points on SSM. These 900 predictions were first reduced to 6 dimensions using PCA, then projected non-linearly into 3 dimensions using Uniform Manifold Approximation and Projection (UMAP), implemented by the Python umap-learn package. The parameters used were ‘n_neighbors’ = 100, ‘min_dist’ = 0.8, ‘metric’ = ‘cosine’, and ‘init’ = ‘spectral’. To visualize the continuous manifolds, we interpolated small surfaces of adjacent x, y coordinate predictions using the plot_surface function in the Matplotlib Python package (Figure 3C).

We visualized other manifold slices similarly. Figure 3E considered four adjacent spatial points (centered at *x* = 75 cm, *y* = 75 cm, with an adjacent points’ distance of 4 cm) and varying speed values. SI Figure 5C considered four distant spatial points (centered at *x* = 75 cm, *y* = 75 cm, with an adjacent distance of 20 cm). SI Figure 5D also considered a slice with a fixed *x* = 75 cm. PCA analysis on these slice manifolds suggested they are low-dimensional (3 PCs are sufficient to explain 80 percent of the variance). Hence, these manifold slices were directly visualized in the space of the first two/three principal components.

### Persistent Homology Barcode

Persistent homology is a method to analyze the topological structure of data clouds^69^. Each point in the data cloud was replaced by a small ball of radius *r*. If the distance of two points is smaller than 2*r*, then they would be connected. Roughly speaking, a graph with dots and connected lines is called a simplicial complex. Simplicial complex can have several holes of different dimensions (0D hole means a single component connecting all points; 1D hole is a loop; 2D hole is a cavity). As the dot ball radius increases from 0 to infinity, different dots would be connected, resulting in different simplicial complexes. During this process, some holes emerge while some holes die out. The birth and dead time of different holes can be collected and represented as bars. All bars of the same hole dimensions form a barcode for that dimension. Usually most bars are short, they are probably noise structure, while long-life bars indicate non-trivial topological structure of the data cloud. The number of long-life bars in each dimension is counted as a Betti number, written in β_*i*_. For example, a loop manifold should have one long-life zero-D hole, one one-D hole and no 2D holes. Hence the corresponding Betti number should be (*β*_0_, *β*_1_, *β*_2_) = (1,1,0). In particular, a torus should have a Betti numbers (*β*_0_, *β*_1_, *β*_2_) = (1,2,1). We used the software package Ripser to compute the barcode, accompanied with approximated sparse filtrations to increase computational efficiency^70^ (epsilon approximation constant = 0.2, see more detail in Ripser^71^).

Intuitively, instead of computing the distance matrix of all points in the data cloud, approximated sparse filtrations discard balls which are completely covered by other balls (under certain *r*).

To build an objective procedure counting the number of long-life bars in the barcode, we defined a bar-length threshold to distinguish long-life bars. Here we defined the length threshold heuristically same as previous study^35^. A data point (e.g. a neural state) is a n-dimensional array where n is the number of grid cells. All data points form a m-by-n matrix, where m is the number of data points. We then randomly rolled (periodic boundary) each column of the matrix. This shuffled dataset was then fed into persistent homology, the maximum bar length was collected. This shuffling procedure repeated 20 times and obtained the final maximum bar length among 20 shuffling. This is the bar-length threshold.

Specifically, 𝒟_*s*_ was used to fit the GKR. When estimating the topological structure of the full three-dimensional manifold, 6,400 random labels were randomly sampled, and input to GKR to generate 6,400 manifold points. To simplify these data points, in align with Gardner et al 2022^35^, we firstly projected these data points into six PC subspace, and then used k-means to compute 1,200 cluster centers. These centers were then fed into Ripser, and Betti numbers were estimated from the above procedure. (*β*_0_, *β*_1_, *β*_2_) = (1,2,1) suggests a successful finding of torus structure in the data cloud.

When estimating the topological structure of SSM (Figure 3C, speed = 20 cm/s), 30-by-30 grid locations were collected, fed into the GKR to make predictions. These 900 manifold points were then projected into the first six PC subspace, and then fed into Ripser to compute Betti k-means approximation in this case, because 900 data points is a good number to computationally handle, unlike the full-manifold case above).

### Computing Geometric Properties of speed-slice manifold at different speeds

𝒟_*s*_ was used to fit the GKR. The fitted manifold is a function of *x, y* locations and speed *v*. For each fixed speed value, we randomly sample 500 locations denoted as (*x*_*i*_, *y*_*i*_).

The SSM center is the averaged 500 manifold points, denoted as ***μ***_*c*_(*v*).

The SSM radius is the averaged Euclidian distance of these 500 random points to the SSM center

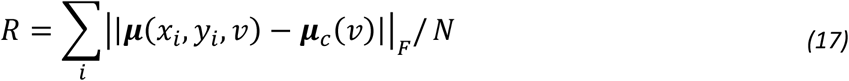

Tangent vector 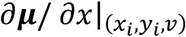 measures how sensitive the neural population representation to the change of *x* location^27^. Let 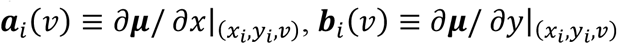 and *a*_*i*_(*v*), *b*_*i*_(*v*) as vector length respectively, lattice area at a point (*x*_*i*_, *y*_*i*_, *v*) is the area formed by two tangent vectors

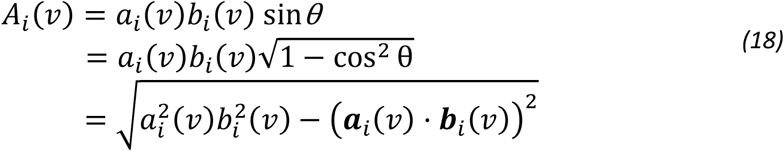

The average of lattice area over 500 random points is the (averaged) lattice area of a SSM, shown in Figure 3F.

Fisher information matrix is defined as *J*^*T*^Σ^−1^*J*, where *J* is the Jacobian matrix in respect to spatial location. Total Fisher information is the trace of Fisher information matrix, averaged over all 500 random points.

To compute the projected noise, Jacobian matrix was normalized to 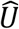 such that each column (tangent vector) has a unit length. Projected noise matrix is

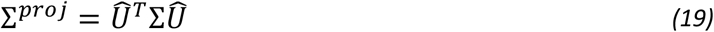

Project noise is the trace of projected noise matrix, averaged across 500 randomly sample points.

### Fisher information provides the upper bound of spatial classification accuracy

Consider a classification problem involving data from two small boxes centered at ***x***_±_ = ***x***_*c*_ ± *δ****x***. Denote the two classes as 𝒞_1_ and 𝒞_2_. In the process of evaluating SCA, we subsampled the data points so that two boxes have equal data set sizes. In line with this, the prior probabilities of a data point belonging to either class are equal, *p*(𝒞_1_) = *p*(𝒞_2_) = 1/2. We also assume the neural state ***r*** in box *i* is approximately given by 𝒩(***r***; ***μ***_*i*_, Σ), where *i* can be 1 or 2. Here we derive the optimal classification accuracy if the classification boundary is linear (as used by the logistic classifier). A linear classification boundary means that the class is 𝒞_1_ if *y* = *w*^*T*^*r* − *w*_0_ < 0, and 𝒞_2_ otherwise.

Classification accuracy is the probability of a correct classification.

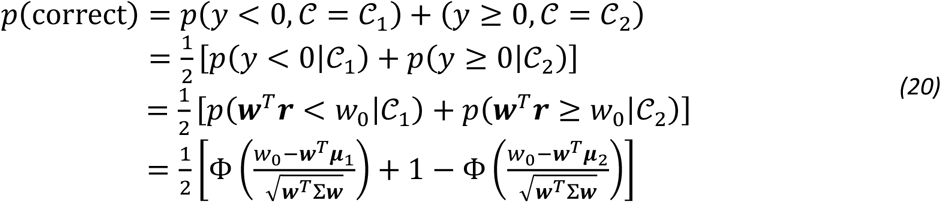

where Φ(·) is the cumulative density function of a standard normal distribution. We used the fact that, if *p*(***r***|𝒞_*i*_) is a Gaussian, ***w***^*T*^***r*** is also a Gaussian with mean ***w***^*T*^***μ***_*i*_ and variance ***w***^*T*^Σ***w***.

Next, we find the optimal *P*(correct). Let ∂*P*(correct)/ ∂*w*_0_ = 0, we get *w*_0_ = ***w***^*T*^(***μ***_1_ + ***μ***_2_)/2. Let ∂*P*(correct)/ ∂***w*** = 0, we get an equation *δ*μ(2***w***^*T*^Σ***w***) = 2Σ***w***(***w***^*T*^*δ****μ***) where *δ****μ*** = ***μ***_2_ − ***μ***_1_. This equation has a general solution ***w*** ∝ Σ^−1^*δ****μ***. Substituting this back into accuracy, we get the optimal accuracy of the two boxes is 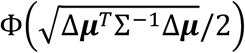.

In our case, boxes were chosen to be symmetric, hence ***μ***_1_ = ***μ***(***x***_*c*_ − *δ****x***) ≈ ***μ***(***x***_*c*_) − ∇***μ****δ****x***, and similarly ***μ***_2_ = ***μ***(***x***_*c*_ + *δ****x***) ≈ μ(***x***_*c*_) + ∇***μ****δ****x***, where *δ****x*** = *δl* **ê**. Therefore, *δ****μ*** = 2∇***μ****δ****x***. The upper bound of accuracy becomes 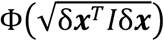 where *I* is the Fisher information. In our numerical procedure, **ê** is a random unit vector. The overall upper bound of accuracy is given by the integration across all angles of **ê**

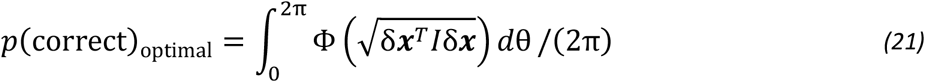

The relationship between optimal accuracy and the total Fisher information becomes intuitive if we assume the Fisher information is unbiased in all directions, which, biologically speaking, means that a rat has no directional bias in an open field. Under this assumption, the integral is trivial because the function inside is independent of direction. Second, the Fisher information becomes proportional to the identity matrix *Tr*(*I*)𝕀. Without loss of generality, let *δ****x*** = *δl*(1,0)^*T*^. The optimal accuracy becomes (Taylor expanded around zero) 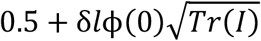 where 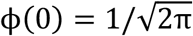. Therefore, optimal accuracy asymptotically increases with the square root of the total Fisher information.

We used Monte Carlo method to estimate the integral in equation (21). Specifically, we sampled 2D vectors from a 2-dimensional standard Gaussian distribution, then rescaled the 2D vectors to have a length equal to *δll*, resulting the sampled *δ****x***. This sampling is unbiased with respect to angle because the 2-dimensional standard Gaussian distribution is isometric. Given Fisher information *I*, the upper bound was estimated as the average of 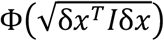 across all *δx*.

### Test upper bound of spatial classification accuracy on synthetic datasets and grid cell population responses

The upper bound (Equation 21) is straightforward in a one-dimensional *δx*

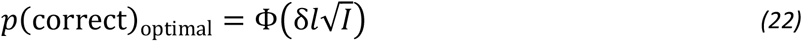

We tested upper bounds in both 1D and 2D synthetic data sets. For each parameter configuration (number of neurons *N*, number of data points *K*, and noise level α; the other parameters were fixed as described in Methods: One-Dimensional and Two-Dimensional Synthetic Datasets), we generated *K* data points. SCA was computed as described in ‘Methods: Spatial classification accuracy’. On the other side, the generated *K* data points were also used for fitting GKR. Fitted GKR make predictions of Fisher information, which then converted to upper bound (Equation 21). Finally, the ground truth Fisher information of the synthetic datasets was also used to compute the upper bound. Results are shown in SI Figure 7.

We also inspected the upper bounds on the Grid cell datasets. GKR provides predictions of Fisher information, which were then used to compute the upper bounds. The upper bounds and SCAs of the R1M2 were shown in SI Figure 6C and Figure 5C for 𝒟_*s*_ and 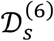 (projection to six PCs, see Methods), respectively. Upper bound-speed/SCA-speed array has 50 × 8 = 400 data points (fifty sampling 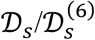 times 8 speed bins). To have a quantitative comparison between upper bounds and SCA, we computed the Pearson correlations between the two arrays, denoted as *r*. The p-value (two-sided) and confidence interval (via Fisher z-transform) can be computed accordingly via python package stats.peasonr^72,73^.

### Independent Firing Grid Cells

We investigated the effect of grid cell activity correlation by comparing results from the original dataset 𝒟_*s*_ to those from hypothetical independent firing grid cells (IFGC). A classic method for generating independent firing cells involves shuffling trials within the same condition. Specifically, each cell’s firing profiles are randomly permuted across trials within a condition. This approach preserves single-cell firing statistics while disrupting cell-to-cell firing correlation. We adapted this method when computing SCA of the IFGC (Figure 6E). Recall that the key idea of SCA is to compute the classification performance on data within two nearby (spatial) boxes. We treat data within each box as a single condition, where each data point (an N-dimensional vector of single-cell firing rates) represents one trial. We then randomly permute each cell’s firing rate across all data points within the box, breaking the cell-to-cell correlation. The SCA of this “trial-shuffled” data is called the SCA of IFGC (Figure 6E).

We also adapted this “trial-shuffling” idea to compute the geometric metrics (total noise, projected noise, and Fisher information) for IFGC. Specifically, after fitting 𝒟_*s*_, GKR can predict mean and covariance at a condition ***x***. Consider GKR as a generative model, it generates infinite data points under the same condition ***x***. If we applied the above “trial-shuffling” procedure on these data points, and recompute the mean and covariance matrix, the mean remains unchanged, while the covariance matrix retains only the diagonal components of the original covariance matrix, with all off-diagonal components set to zero. Therefore, IFGC’s GKR is same as the original GKR except only having the diagonal covariance matrix. With IFGC’s GKR, the geometric metrics can be computed as previously described.

We compared speed-averaged metrics obtained from the original datasets to that obtained from IFGC. Methods of computing speed-averaged metrics along with statistical analysis can be found in the Methods: Bayesian linear ensemble averaging and statistical testing section.

## Author contributions

Conceptualization: ZY, RW Methodology: ZY, RW Investigation: ZY Supervision: RW

Writing: ZY, RW

## Data and materials availability

The analysis code is available at https://github.com/AgeYY/speed_grid_cell_information.git

## Supplementary Information

### SI Methods: Mathematical details of Bayesian Linear Regression and statistical testing

First, we consider only one sampled dataset 𝒟_*s*_. From this dataset, we obtain a metric-speed dataset (e.g. SCA-speed in Figure 1C or SSM Radius-speed in Figure 3G), denoted as {*t*_*i*_, ***x***_*i*_}, where *t*_*i*_ is the metric value and ***x***_***i***_ = (*v*_*i*_, 1) with *v*_*i*_ denoting the speed. Assuming

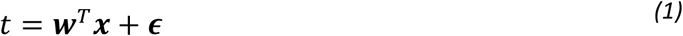

where ***ϵ*** ∼ 𝒩(***ϵ***| 0, *β*_*t,s*_^−1^) *β*_*t,s*_ is a scalar representing precision. Bayesian Linear Regression (BLR) is used to obtain the posterior distribution *p*(***w***|𝒟_*s*_), which is a Gaussian distribution 𝒩***w***; *m*_*w,s*_, Σ_*w,s*_. Substituting this back to (1), we obtained the predictive distribution p(*t*_*q*_***x***_*q*_, 𝒟_*s*_) as 𝒩*t*(_*q*_; *m*_*t,s*_, Σ_*t,s*_), where 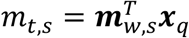 and 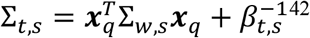

Next, we consider the whole dataset 𝒟, taking into account all 𝒟_*s*_. Each 𝒟_*s*_ is a random subsampling of 𝒟, therefore, *p*(*w*|𝒟) = ∑_*s*_ *p*(*w*|𝒟_*s*_)*p*(𝒟_*s*_ |𝒟) = ∑_*s*_ *p*(*w*|𝒟_*s*_)/ *B*, where *B* = 50 is the number of samplings. This distribution is a mixture of the Gaussian, we approximated it as a single Gaussian function with the same mean and covariance. The mean of *p*(*w*|𝒟) is

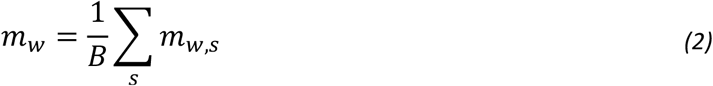

The covariance is

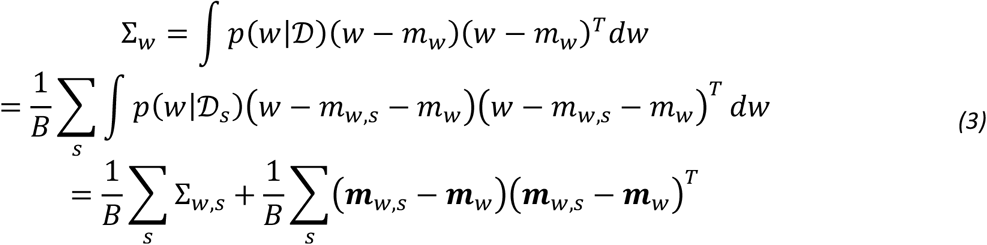

We can use the same trick to compute the mean and covariance of the predictive distribution. The mean is

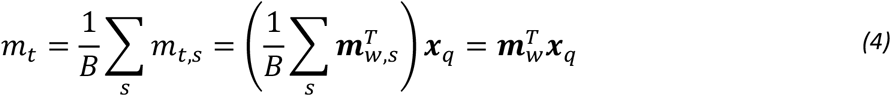

The covariance is

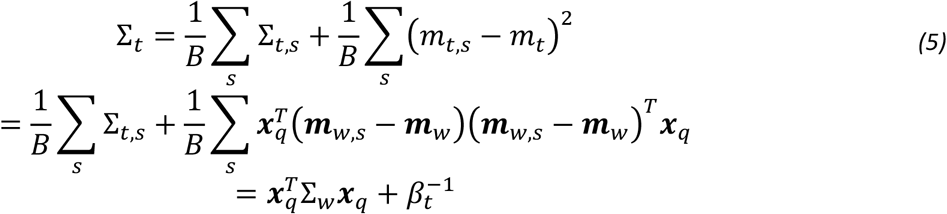

where 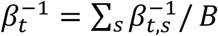. Inspecting the mean and covariance of the predictive distribution, it is clear that even considering the whole dataset 𝒟, metric is still a linear function of speed, written explicitly as

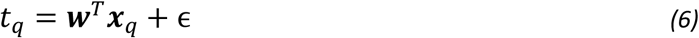

where ***w*** ∼ 𝒩(***w***; ***m***_*w*_, Σ_*w*_) and 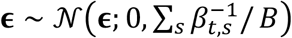

**SI Figure 1.**
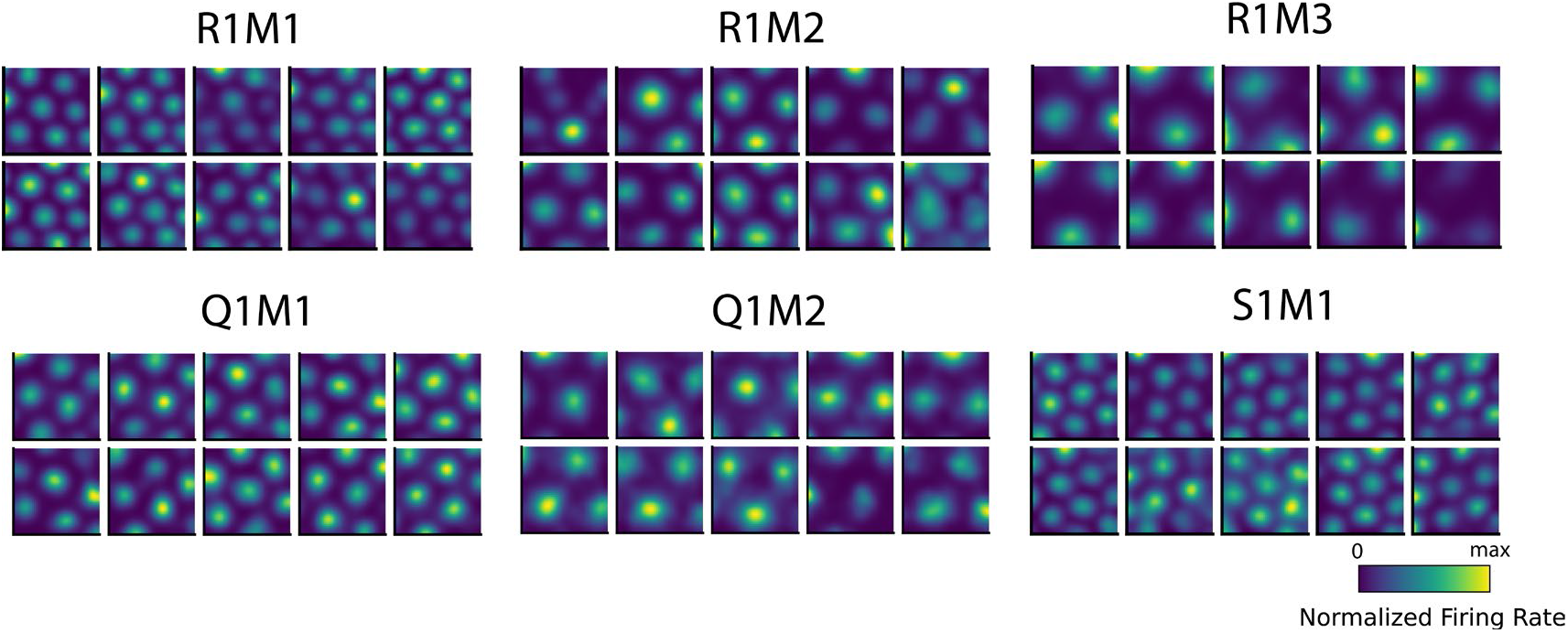
Example grid cells’ rate maps from different datasets (see Methods).

**SI Figure 2.**
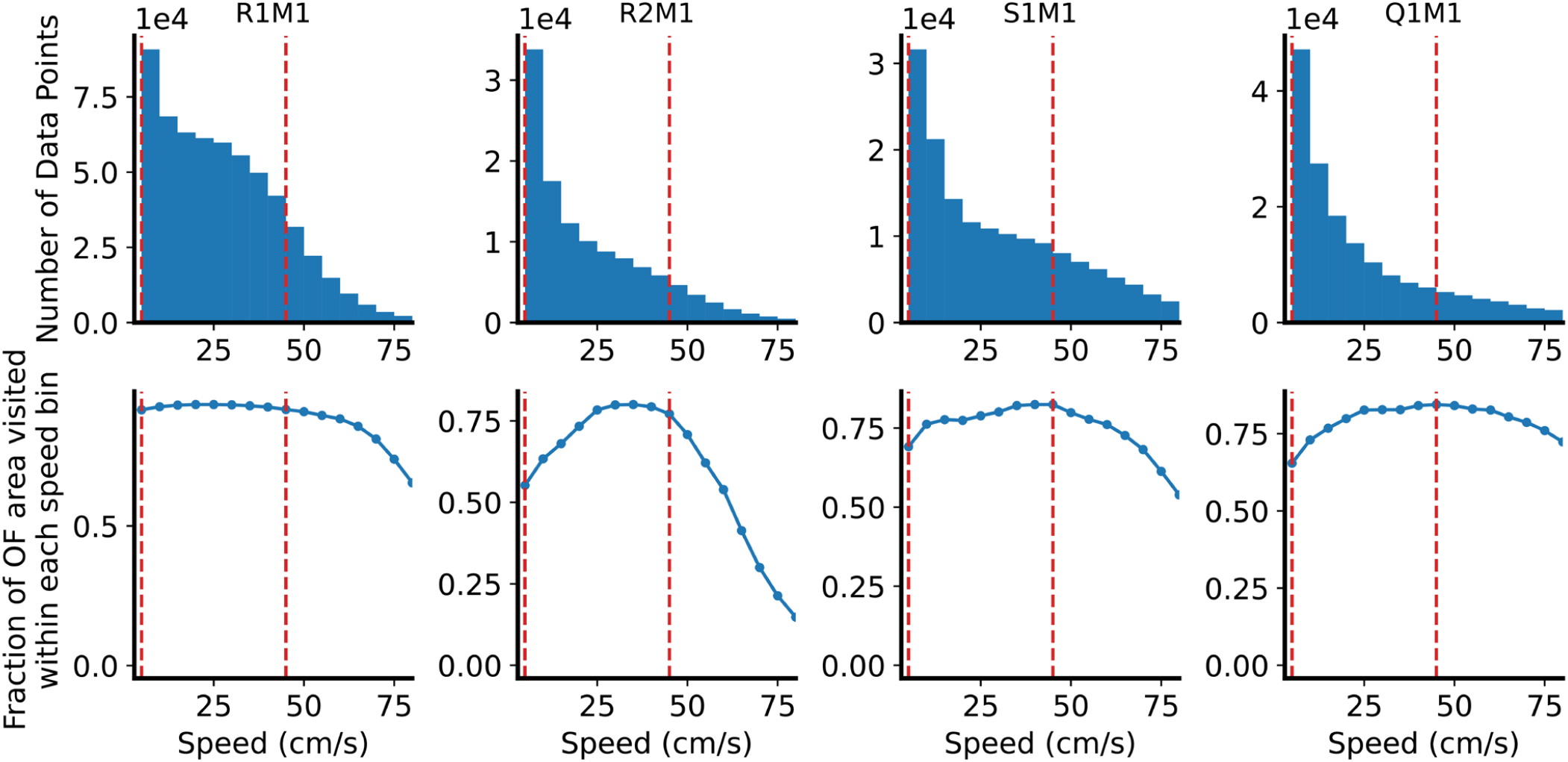
Statistics of behavior labels (x, y locations and speed). Upper: Number of data points in each small speed bin (bin width = 5 cm/s). Each data point represents a neural state at a 10 ms time bin. Two vertical dashed lines enclose the speed range considered in this paper (5 cm/s to 45 cm/s). The statistics for R1M2 and R1M3 are the same as R1M1; R2M2 and R2M3 are the same as R2M1; Q1M2 is the same as Q1M1. Bottom: The entire OF area is digitized into 30-by-30 spatial bins. The y-axis indicates the fraction of bins visited by the rat within a speed bin.

**SI Figure 3:**
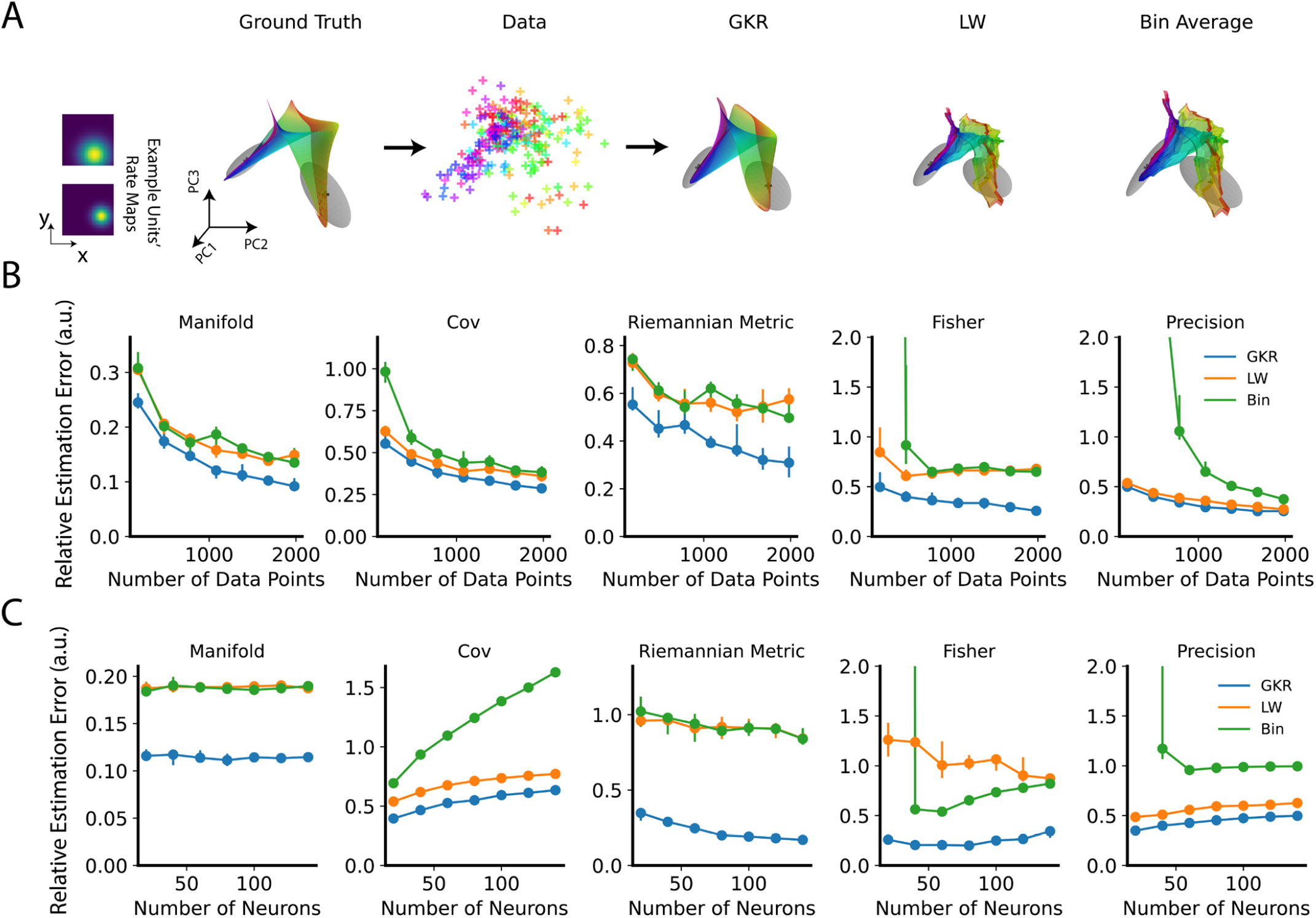
Testing GKR on a 2D synthetic manifold. (**A**) The synthetic dataset comprises *N* synthetic neurons with heterogeneous tuning maps on a 2D space ***P***. Ground truth ***μ***(***P***) and Σ(***P***) were visualized using the first three principal components, shown on the left. Ellipsoid axes represent the directions of three covariance eigenvectors, with lengths proportional to the eigenvalues. In this example, the synthetic dataset had 10 neurons and generated 200 data points. These data points were then fed into different methods for manifold inference. (**B, C**) Evaluation of different methods’ performance under various conditions. The default number of data points is 1,000, and the default number of neurons is 10. The illustration of these panels is the same as in main Figure 2C, D.

**SI Figure 4.**
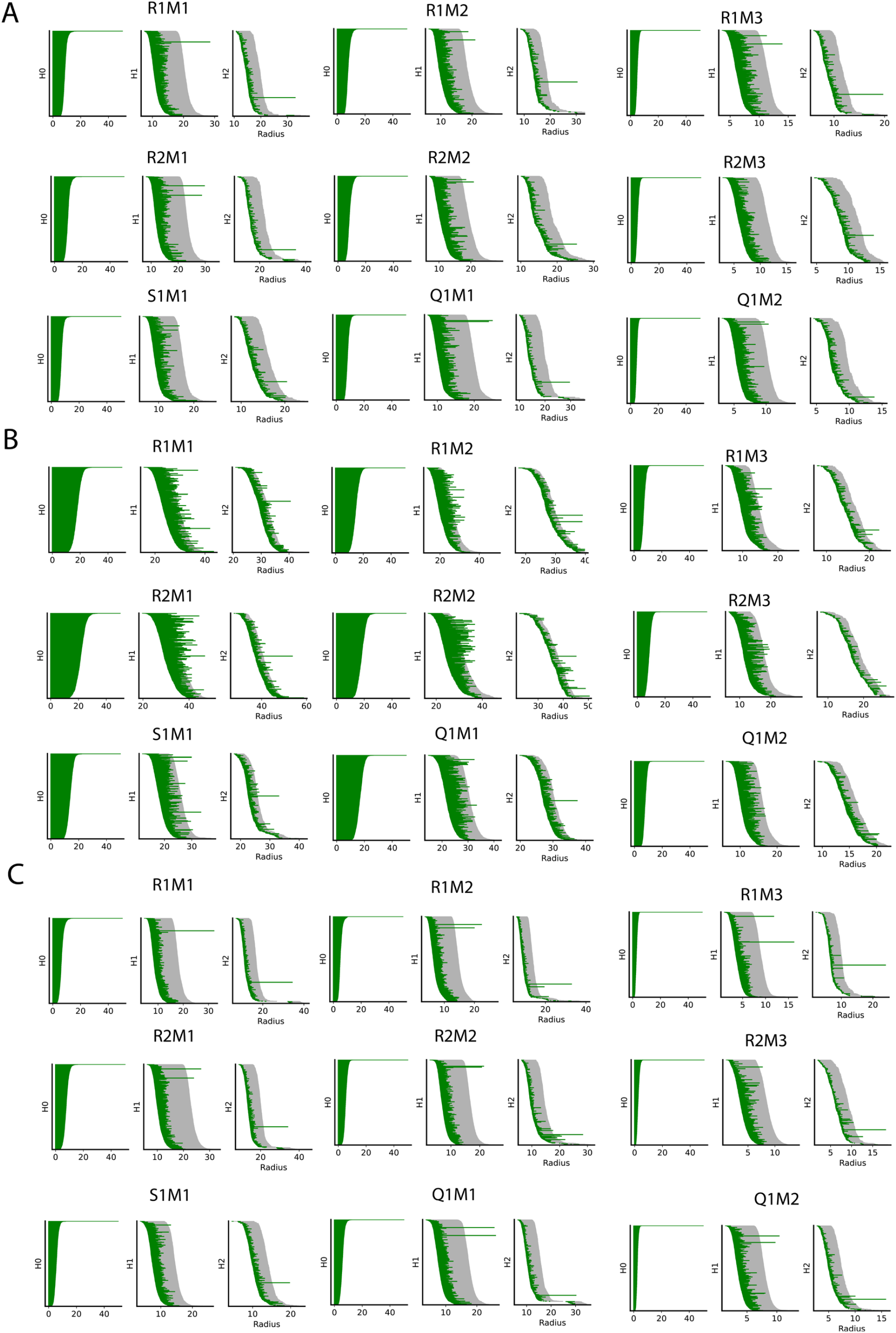
Grid cell population forms toroidal-like manifolds. (**A**) Persistent homology barcode for topological analysis. Long bars represent possible true topological structures. H0, H1, and H2 indicate a connected component, a circular hole, and a cavity, respectively. A torus is characterized by one long bar in H0, two in H1, and one in H2. The sampled dataset 𝒟_*s*_ was used to fit a GKR model. The fitted manifold is intrinsically three-dimensional (with three labels: x location, y location, and speed). We randomly sampled 6,400 label points and input them into the GKR model to predict 6,400 manifold points in the original high-dimensional space (where the number of dimensions equals the number of grid cells). These manifold points were then reduced to their first six principal component (PC) dimensions. These dimensionally reduced manifold points were clustered into 1,200 centers using k-means clustering. These 1,200 cluster centers were then analyzed using persistent homology, as shown by the barcode in the figure. Grey bars indicate the maximum bar lengths from 20 shuffles of the 1,200 cluster centers (see Methods). (**B**) Same as (A), but without PCA dimension reduction. (**C**) Similar to (A), but with speed fixed at 20 cm/s. At this speed, 30-by-30 grid points were sampled in the OF space, fed into GKR to predict 900 manifold points, which were then projected onto the first six PC dimensions and analyzed using persistent homology (see Methods). Grey bars indicate the maximum bar lengths from 20 shuffles of the 900 manifold points (see Methods).

**SI Figure 5.**
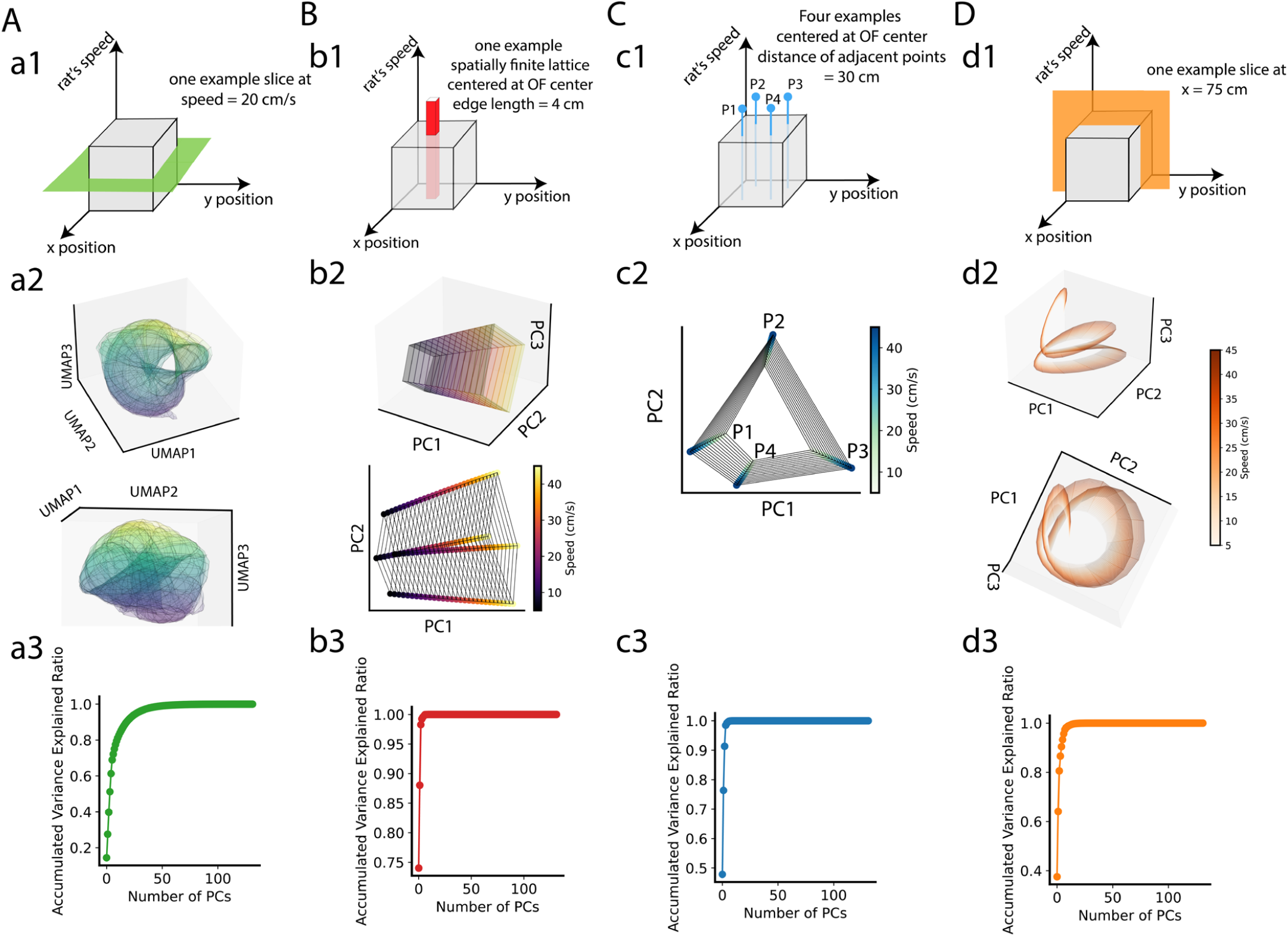
Visualization of different manifold slices of a GKR fitted from R1M2. (**A**) a1: The speed slice; a2: Two views of the manifold, visualized by first projecting the manifold to the 6 PC space, then non-linearly reduced to three dimensions using UMAP; a3: Accumulated variance explained ratio of the manifold. (**B, C, D**) Same as (A), but for different slices. Due to their low dimensionality, manifold slices were directly visualized in the PC spaces.

**SI Figure 6.**
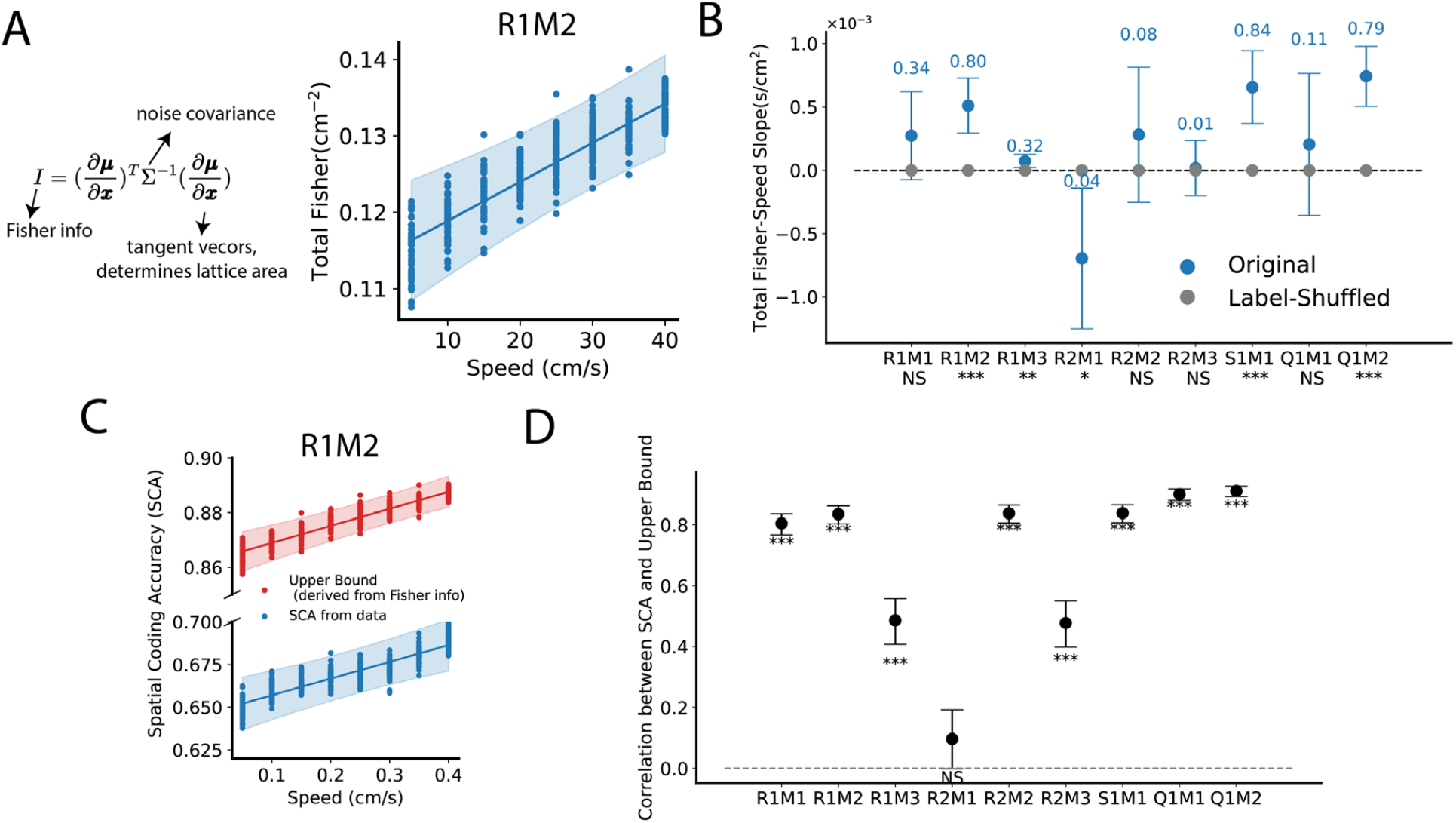
Speed modulation of Fisher information computed from the original high-dimensional space (dimensionality equals to the number of grid cells). This figure is same as Figure 5, but using the original 𝒟_*s*_ without PCA reduction.

**SI Figure 7.**
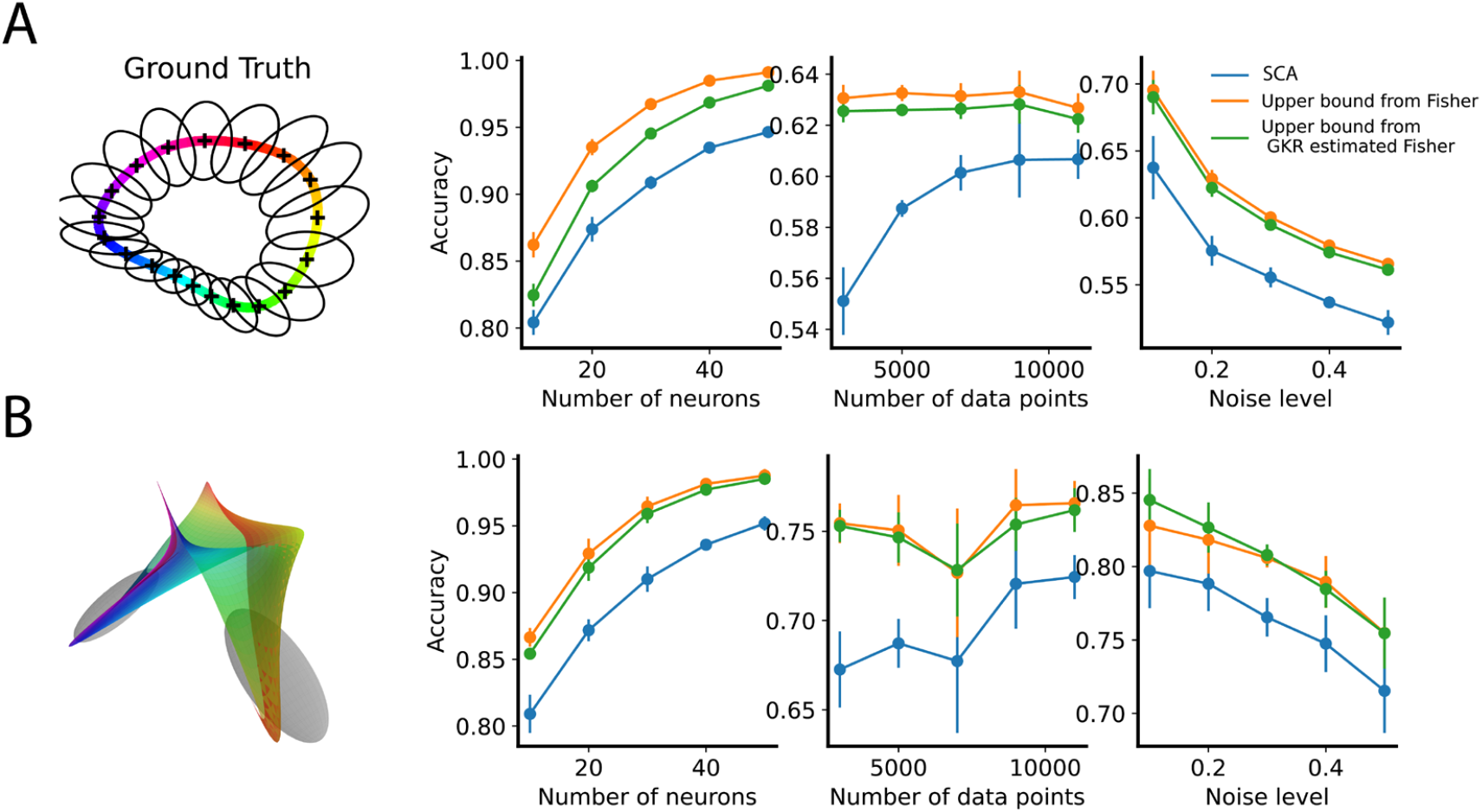
Testing upper bounds of the SCA derived from Fisher information on synthetic datasets. (**A**) The default parameters are 5 neurons, 5000 data points, and noise level ν = 0.2. Other parameters are detailed in Methods. For each condition, data points were input into GKR, which outputted estimated Fisher information. GKR’s estimated Fisher information was then used to compute the SCA upper bound (see Methods). We also computed the upper bound using ground truth Fisher and directly calculated SCA from the raw data points (see Methods). Dots and error bars represent the median, first, and third quantiles from 10 samplings. (**B**) Same as (A) but using 2D synthetic datasets. The default parameters are 5 neurons, 10,000 data points, and noise level ν = 0.5. Other parameters are detailed in Methods.

